# Diversity, connectivity and negative interactions define robust microbiome networks across land, stream, and sea

**DOI:** 10.1101/2025.01.07.631746

**Authors:** Kacie T. Kajihara, Mengting Yuan, Anthony S. Amend, Nicolas Cetraro, John L. Darcy, Kauaoa M.S. Fraiola, Kiana Frank, Margaret McFall-Ngai, Matthew C.I. Medeiros, Kirsten K. Nakayama, Craig E. Nelson, Randi L. Rollins, Wesley J. Sparagon, Sean O. I. Swift, Mélisandre A. Téfit, Joanne Y. Yew, Danyel Yogi, Nicole A. Hynson

## Abstract

In this era of rapid global change, factors influencing the stability of ecosystems and their functions have come into the spotlight. For decades the relationship between stability and complexity has been investigated in modeled and empirical systems, yet results remain largely context dependent. To overcome this we leverage a multiscale inventory of fungi and bacteria ranging from single sites along an environmental gradient, to habitats inclusive of land, sea and stream, to an entire watershed. We use networks to assess the relationship between microbiome complexity and robustness and identify fundamental principles of stability. We demonstrate that while some facets of complexity are positively associated with robustness, others are not. Beyond positive biodiversity *x* robustness relationships we find that the number of “gatekeeper” species or those that are highly connected and central within their networks, and the proportion of predicted negative interactions are universal indicators of robust microbiomes. With the potential promise of microbiome engineering to address global challenges ranging from human to ecosystem health we identify properties of microbiomes for future experimental studies that may enhance their stability. We emphasize that features beyond biodiversity and additional characteristics beyond stability such as adaptability should be considered in these efforts.

## Introduction

Stability, or the ability of biological communities to maintain their functions in the face of change, is paramount for the persistence of ecosystem services upon which all life on our planet relies. For over 70 years ecologists have debated the relationship between stability and ecosystem complexity (the biodiversity of an ecosystem and the interactions therein^1,2^). Counter to previous paradigms put forth by Odum (1953)^3^, Elton (1958)^4^ and others, supporting a positive complexity *x* stability relationship, May’s 1972^5^ seminal paper proposed that more complex communities should be less stable. Among ecologists a resounding critique of May’s work was that these mathematical models did not represent real world systems. In response, food webs emerged as model natural study systems to determine the principles of stability^6^. However, even from these decades-long efforts, food web ecologists have yet to agree on the relationship between stability and complexity. Some propose complexity, inclusive of factors such as richness^7^, trophic interactions^8^ and phylogenetic diversity within and among guilds^9^, among others, is a fundamental property of stability as it buffers against extinction cascades; while others propose the opposite, that complexity potentially reduces the proportion of strong interactions among species leaving food webs more susceptible to collapse^1,2,10^. More recently it has been predicted that other facets of complexity such as dominantly competitive, or other negative interactions among species should enhance stability by diffusing the spread of perturbations, while others find that instead, mutualisms are stabilizing^2^. Part of the incongruence among studies may be that the definition of complexity varies among studies, and study systems vary in their innate complexity. Therefore, identifying inherently complex systems and measuring multiple facets of complexity to examine stability *x* complexity interactions may help reconcile some of these differences. Microbiomes offer this opportunity as they are some of the most complex biological communities on earth often involving interactions among thousands of taxa and spanning the spectrum of biotic interactions and inhabiting basically every organism and environment on the planet^11^.

While microbiomes may not always partition into discrete guilds like food webs, properties affecting their stability and robustness may be similar^12,13^. In ecological communities macro-organisms engage in complex relationships with each other ranging from positive (e.g., mutualism, commensalism) to negative (e.g., parasitism, amensalism), which together influence community composition and the health of hosts and ecosystems^14^. Microorganisms are no different^15^, taking part in intricate webs of interactions with other microbes, hosts, and the environment that sustain the metabolic and biogeochemical backdrop against which life persists^16,17^. Enhancing microbiome stability has recently come under the spotlight as an aspirational goal for management and engineering efforts, to encourage microbiomes to successfully establish and maintain their functions^18,19^. Stability is also considered a key factor for microbiomes to resist or remain resilient against disturbances such as climate change, changes in host diet, or antibiotic treatments that can profoundly affect diversity and community composition and lead to alternative stable states which may, or may not be desirable^20,21^.

Stability is a property influenced by many interacting factors including community resistance and resilience to disturbance, as well as how a community responds to species losses^22^. Extinction cascades or, the degree to which the loss of one species impacts the loss of others in the same system, generates variation in community robustness, which is an important measure of stability^23^. In the case of microbiomes, extinction cascades can not only lead to a loss of microbial biodiversity, but also potentially profoundly affect host and ecosystem function^13^. Currently, it is unclear whether there are universal principles that govern microbiome stability, or whether certain microbiomes are more robust than others. While some hosts and environments harbor specific microbes, many microbes traverse these boundaries and microbiomes in general have a tendency towards nestedness^24^. Therefore, assessing the guiding principles of microbiome stability demands an ecosystem-scale approach.

Unlike many macro-organismal food webs, it is challenging to directly observe phenomena such as extinction cascades, competition, or keystone species in microbiomes due to their complex, ephemeral, and microscopic nature. Methods to overcome these challenges include computational tools such as co-occurrence networks built from targeted or untargeted metagenomic data^25^, which are a practical lens to assess various components of stability including robustness. In these networks adopted from graph theory, nodes represent taxa and edges represent statistically significant occurrence or abundance associations between them, either positive or negative. An important caveat for these computational methods is that they generate predictions of biologically meaningful interactions, which need to be validated. However, the power of network methods lies in their ability to embrace the often otherwise intractable diversity of microbiomes to generate strong scalable hypotheses, which can then be tested through more reductionist approaches. Furthermore, co-occurrence patterns (e.g. presence/absence) form the fundamentals of community assembly regardless of whether members directly interact or not^26^.

Similar to stable food webs, stable networks should be robust to node removal^27^, meaning their structures resist rapid collapse when nodes are removed. However, not all nodes are “created equal” and similar to the concept of keystone species, the removal of highly connected and central nodes within a network should lead to more rapid collapse^28^. Other node-specific or global network properties related to complexity should also impact robustness. For example, node richness and the ratio of edges to nodes should be positively correlated with robustness if there is a positive complexity *x* stability relationship. Modularity, which measures the partitioning of species into distinct and highly connected sub-communities, should also have a positive relationship with network stability where higher modularity should contain the effects of disturbance to specific modules rather than impacting the entire network^29^, a concept reminiscent of what May^5^ referred to as “blocks.” Another is connectance or the number of realized edges in a network among all the possible ones. Higher degrees of connectance within a network should result in greater robustness due to more paths among nodes damping the effect of changes in any one node’s persistence on the persistence of others, similar to the concepts derived from MacArthur’s 1955^30^ models of population and community stability. However, whether these, or other network properties universally increase microbiome robustness has yet to be established.

We broadly define microbiome robustness as the relative ability to maintain network structure in the face of node removal. We predict that the sequential removal of highly central and connected nodes from networks will have the greatest negative effect on robustness relative to the removal of less central and less connected nodes, with the effect of random node removal intermediate between these two extremes. We also predict that network robustness will be positively correlated with multiple measures of complexity including richness, connectance, edge to node ratio, predicted negative interactions, modularity and phylogenetic diversity.

We define a universal feature of microbiome robustness as a property that consistently predicts robust microbiomes across networks regardless of the spatial scale they represent. To determine whether there are universal properties of robust microbial networks requires a tractable, yet diverse study system, and so far studies are confined to single systems and sample types (mostly soil), and often single domains of microbial life (mostly bacteria), limiting their generalizability^31^. We address this by capturing free-living and host-associated microbiome diversity across a remarkably steep environmental gradient including connected marine, freshwater stream, and terrestrial habitats in a spatially compact and experimentally tractable watershed in Waimea, Oʻahu, USA (Figure S1). Our model ridge-to-reef Hawaiian ecosystem spans an entire hydrologic cycle and four Köppen climate types, thus our study system plausibly reflects microbial diversity and dynamics at much broader geographic scales^24^.

## Results

### Microbiome networks are non-random

We generated 33 networks representing fungi, bacteria and interkingdom co-occurrences for an entire watershed, its constituent marine, stream and terrestrial habitats, as well as sites along a steep environmental gradient within the watershed (Figs. 1, S2-S7, Table S1). Spatial autocorrelation of operational taxonomic unit (OTU) abundances was not significant in fungi (r = 0.018, P = 0.072), and was significant, but weak, in bacteria (r = 0.055, P = 0.001). Fungal and bacterial OTUs co-occurred across hosts and environmental substrates (Fig. S8), and module composition indicates that network interactions and properties are not constrained by specific hosts or environmental substrates (Fig. 1d-f). Networks exhibited non-random interactions, as determined by their non-Poisson degree distributions and small world properties (Tables S2). Network node degree distributions generally fit a power-law function with a few exceptions: watershed-wide bacterial and interkingdom node degree distributions both fit best to a gamma distribution^32^, and interkingdom node degree distributions in the stream and terrestrial networks, as well as the terrestrial bacterial network all fit best to the Weibull distribution^33^ (Table S2). For these networks, as additional tests of non-randomness, we compared their clustering coefficients (*Cl*, the average proportion of pairs of nodes one edge away from a node that is also linked to each other) and connectance (*C*), which for a random network should be equal^34^. In all cases *Cl* was not equal to *C* (Table S3). The relationships between node degree and node betweenness centrality were significant and positively correlated for all networks (R^2^ 0.331-0.633, P<0.001, Fig. S9).

**Figure 1.**
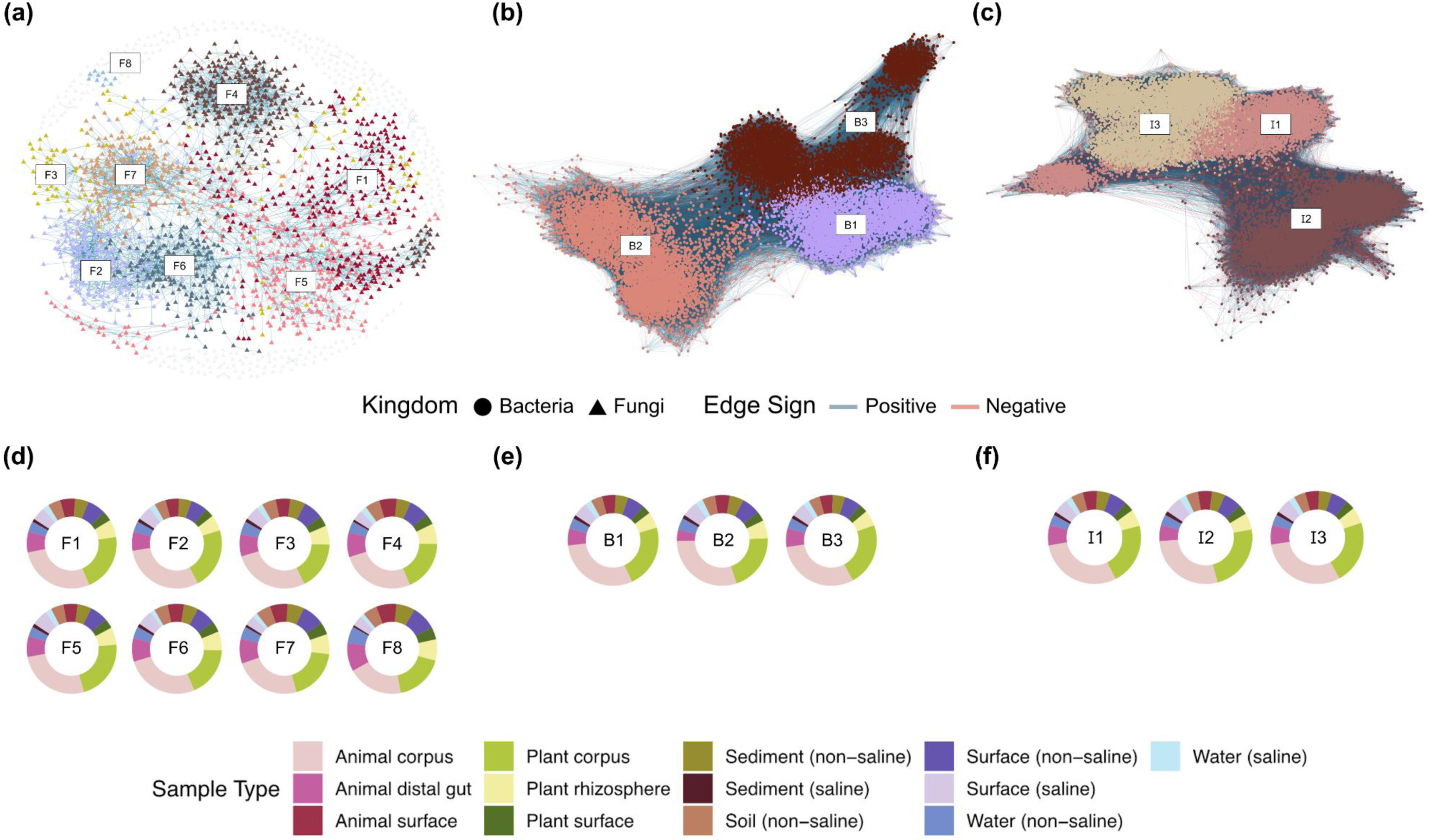
Microbial co-occurrence networks representing an entire watershed. a-c, Visualizations of fungal (a), bacterial (b), and interkingdom (c) networks. Networks are colored and labeled by module, with fungal modules beginning in “F”, bacterial modules beginning in “B”, and interkingdom modules beginning in “I”. For visual clarity, only fungal modules with more than 10 Operational Taxonomic Units (OTUs) are shown. Node shapes are delineated by microbial kingdom and edge colors by edge sign (positive or negative). d-f, Pie charts representing the host and environmental substrate associations of each module. Pie chart sections correspond to the percentages of samples harboring module OTUs that originated from a given host or environmental substrate.

### Interkingdom networks harbor more connected and centralized taxa than bacterial or fungal alone

The number of connections among taxa or node degree, was highest among interkingdom networks, followed by bacteria and then fungi for the entire watershed (P<0.001), habitats (P<0.001), and all gradient sites (P<0.001; Figs S10-S12, Table S4). Similarly, across the entire watershed, habitats and the gradient, interkingdom betweenness centrality values were also significantly higher than those of fungi and bacteria (P<0.001, Figs. S10-S12, Tables S5). This indicates that bacterial and fungal nodes became more connected and more centralized in the networks when considered together rather than independently. In the watershed, fungal and bacterial networks’ betweenness centrality values were not significantly different from each other (P=0.940, Table S5); whereas among the habitats and gradient sites betweenness centrality was significantly higher for fungal networks than for bacterial (P≤0.033, Figs S10-S12., Table S5). Within domains, unique patterns of node degree and betweenness centrality values emerged among habitats and across the gradient sites, with network properties for interkingdom generally more similar to those of bacteria than fungi (Figs S10-S12, Tables S6-S7)

### Universal principles of microbiome network stability

We identify universal properties of robust microbiome networks as those with highly centralized and connected taxa, and those dominated by predicted negative interactions. We find these patterns to hold across all spatial scales from the entire watershed, to its constituent marine, stream and terrestrial habitats as well as along a strong terrestrial environmental gradient (Figs. 2 & 3, S13). In each network 84%-100% (average 97.95% SD±3.63%) of the taxa were encompassed within the starting largest connected component. At all scales including the watershed (Fig. 2a), habitats (Fig. 2b-d), and the gradient sites (Fig 2e-k), the removal of taxa with high betweenness centrality led to more rapid decay of network structure relative to the removal of taxa with low betweenness centrality or at random. Contrary to our prediction, the removal of nodes with low betweenness centrality or random removal generally produced similar patterns of robustness, except in the case of random node removal in some fungal networks that led to more rapid network collapse (Fig. 2, Fig. 4, Fig. S14). Interkingdom networks involving fungal and bacterial co-occurrences were more stable than fungal networks alone, and predominantly more stable than bacterial networks alone (Fig. 2, Table S8).

**Figure 2.**
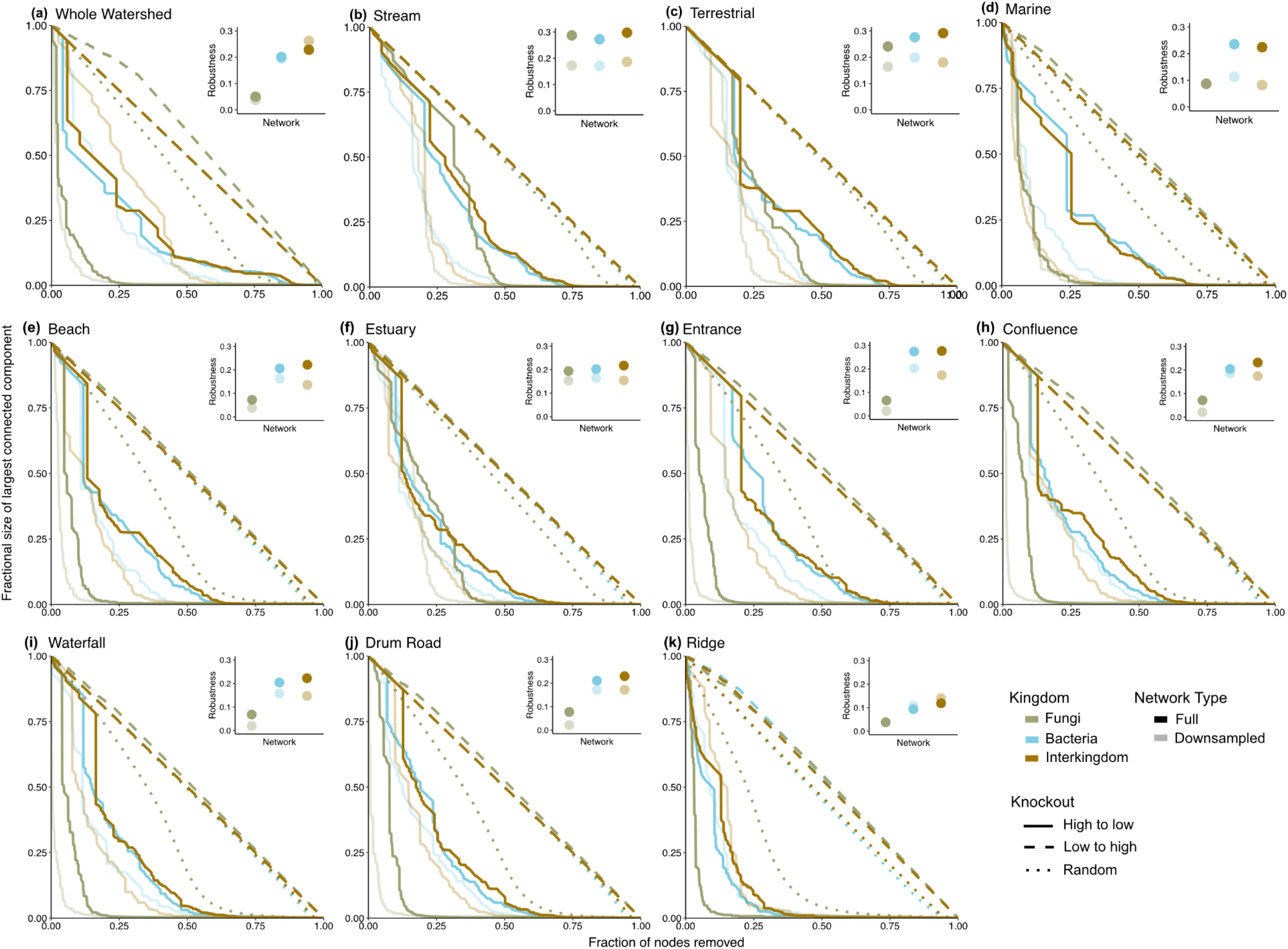
Removal of taxa with high betweenness centrality leads to more rapid network collapse than removal of those with low betweenness centrality or random removal. Attack robustness of microbial co-occurrence networks representing the watershed (a), habitats within the watershed (b-d) and sites along a steep environmental gradient within the watershed (e-k). Robustness is measured as the size of the largest remaining network component relative to its starting size (which in this case included an average of 97.95% SD±3.63% of all nodes) after nodes are removed in order of high betweenness centrality (dark solid lines), low betweenness centrality (dashed lines), or at random (dotted lines). The lightened lines on each panel represent removal of nodes with high betweenness centrality from downsampled networks each with the same number of nodes as the smallest network (721). Each line represents a network from either fungi (green), bacteria (blue) or interkingdoms (brown). More robust networks are indicated by a larger area under the curve. The dots in each subpanel represent each networks’ robustness metric as measured by area under the curve.

**Figure 3.**
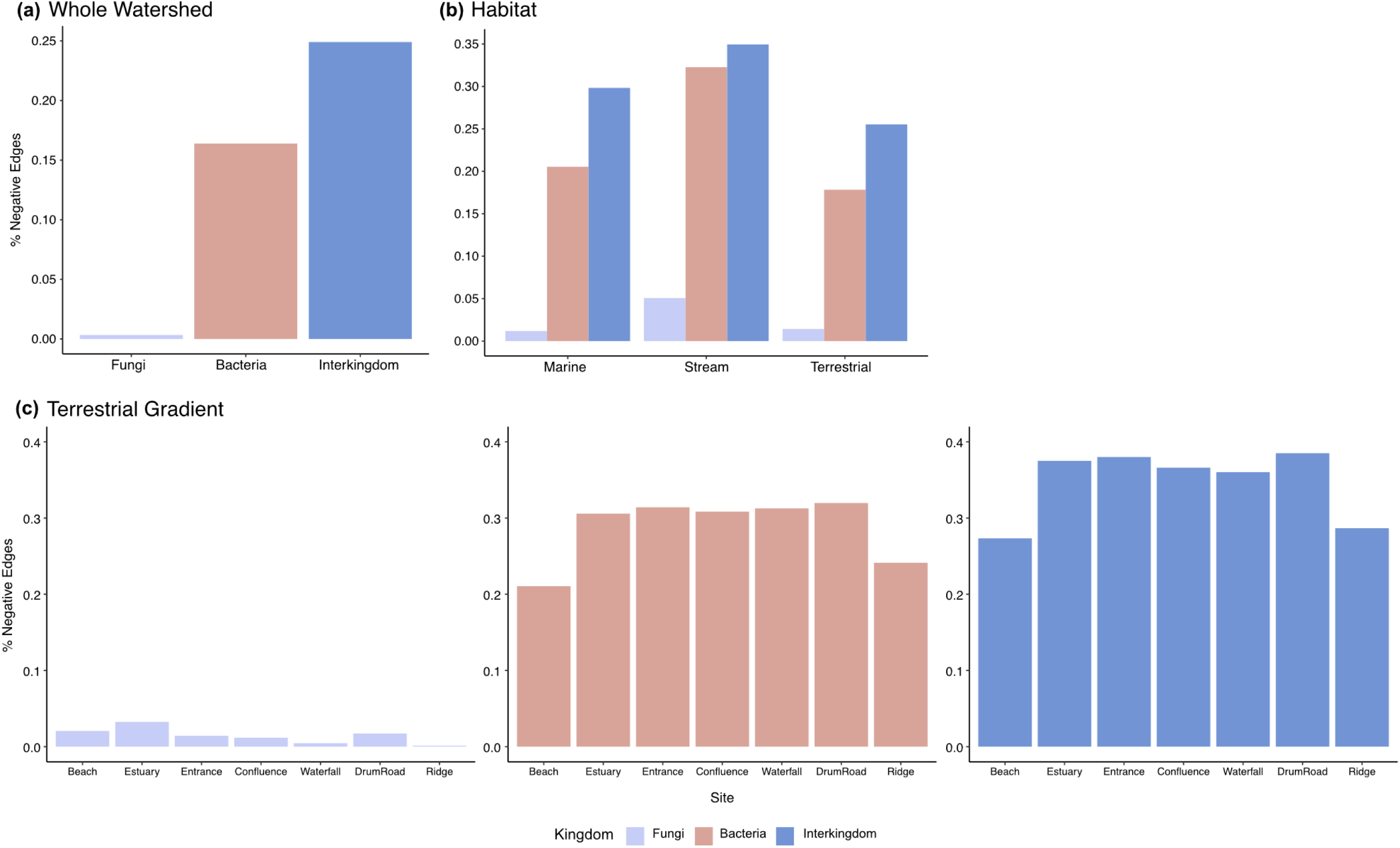
Interkingdom networks consistently have a greater proportion of negative edges in every respective network type. Proportion of negative edges in whole watershed networks (a), habitat networks (b), and gradient networks (c; left, right, and center). All bars are colored by kingdom. For gradient networks (c), sites are listed from left to right in order from the mouth to the headwaters of the watershed.

**Figure 4.**
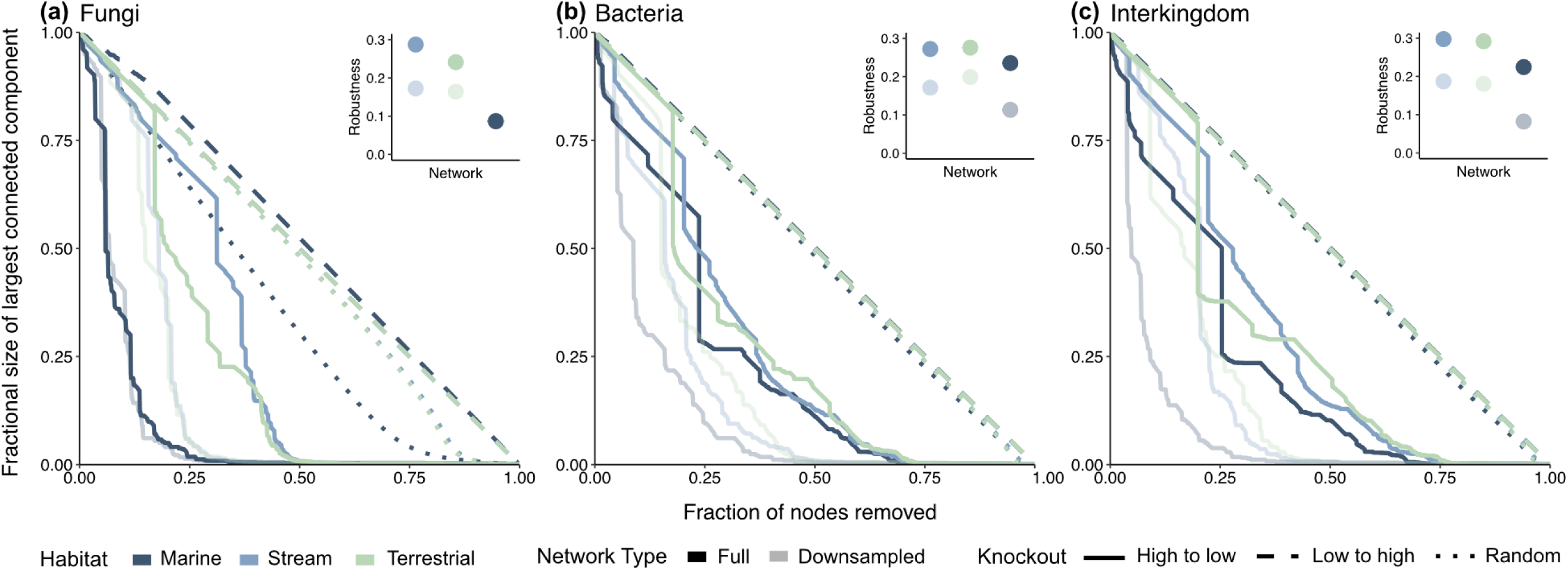
The removal of taxa with high betweenness centrality leads to rapid network collapse especially in marine habitats across fungal, bacterial and interkingdom networks. Attack robustness of fungal, bacterial, and interkingdom networks by habitat (a-c). Robustness is measured as the size of the largest remaining network component relative to its starting size (which in this case included an average of 99.35% SD±1.53% of all nodes) after nodes are removed in order of high betweenness centrality (dark solid lines), low betweenness centrality (dashed lines), or at random (dotted lines). The lightened lines on each panel represent removal of nodes with high betweenness centrality from downsampled networks each with the same number of nodes as the smallest network (721). Each line represents either the marine (dark blue), stream (light blue) or terrestrial (green) habitat. More robust networks are indicated by a larger area under the curve. The dots in each subpanel represent each networks’ robustness metric as measured by area under the curve.

Marine networks were less robust than stream or terrestrial ones (Fig. 4) and marine bacteria were significantly more closely related to one another than in the other two habitats (P < 0.001; Fig S13, Table S8 & S9). Across fungal, bacterial and interkingdom networks the ridge site at the headwaters of the watershed was the least robust among other gradient networks, and bacteria there were also significantly more phylogenetically clustered (P ≤ 0.007; Fig. S13 & S14, Tables S8 & S10). The relationship between phylogenetic diversity and habitats or sites was not biased by sample richness (R^2^ = 0.006, P < 0.001, R^2^ = 0.006, P < 0.001, respectively; Fig. S15).

Overall we found a positive robustness *x* complexity relationship, but not every measure of complexity was positively correlated with robustness. Network robustness was correlated with OTU richness (P=0.001, R^2^=0.30, Fig. 5), but when we controlled for richness by downsampling each network to equal numbers of fungal, bacterial and interkingdom nodes, consistent additional predictors of robustness remained (Fig. S16). As predicted, connectance (R^2^=0.53; P<0.001), edge to node ratio (R^2^ =0.32; P<0.001), and proportion of predicted negative interactions (R^2^ =0.42; P<0.001), were significantly and positively correlated with observed network robustness (Fig. 5). However, counter to our prediction, we found a strongly significant and negative correlation between robustness and modularity (P<0.001, R^2^=0.84). When controlling for differences in node richness, robustness remained similar across the watershed networks, but habitat networks began to collapse more rapidly (Fig. 2a-d). Specifically, the robustness of the marine interkingdom network decreased and became more similar to that of marine fungi, which were largely unaffected (Fig. 2d & Fig. 4). Overall patterns of robustness largely remained consistent regardless of whether richness was held constant or not - the marine habitat and ridge site were the least robust as were fungal networks relative to bacterial and interkingdom (Fig. 2, Fig. 4 & Fig. S14). By leveraging the global-scale heterogeneity of our study site we posit that these properties are not context dependent, but rather potential fundamental rules of life for microbial interactions.

**Figure 5.**
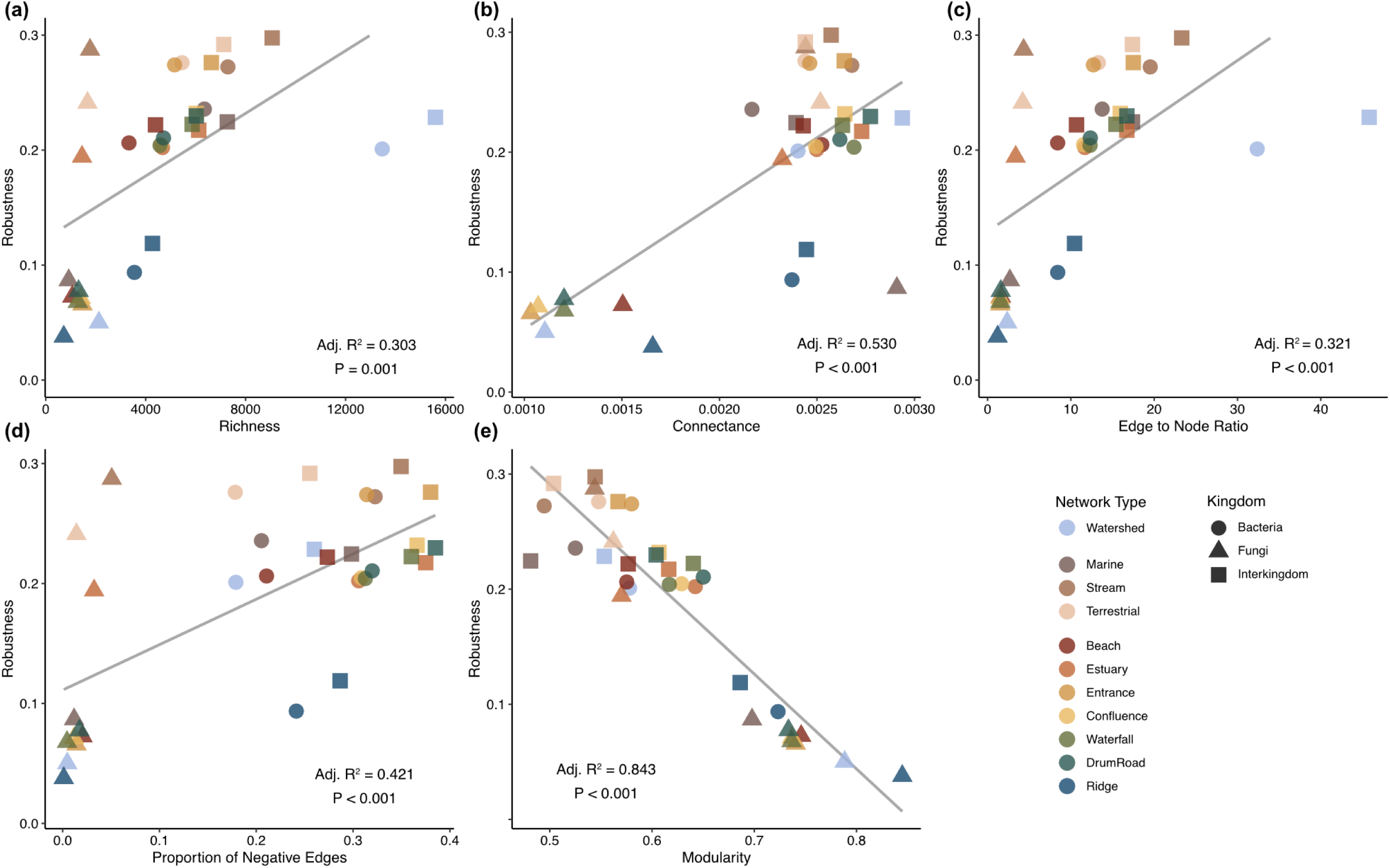
Most measures of complexity are positively related to network robustness except modularity. Linear regressions examining the relationship between robustness (area under the curve from figure 2) and various measures of complexity including richness (a), connectance (b), edge to node ratio (c), proportion of negative edges (d) and modularity (e). Each symbol represents either bacterial (circle), fungal (triangle) or interkingdom networks (squares) and each color a network of different spatial scale. All relationships are statistically significant at α≤0.05 and p<0.001 to p=0.001.

## Discussion

The stability of networks ranging from ecosystems to the internet to neuron pathways in the brain, or in this case, microbiomes, is affected by numerous properties of these systems inclusive of resistance and resilience against disturbance^8,35^, the ability to “rewire” interactions^36^, as well as their robustness or the ability to maintain structure and function in the face of loss such as brain damage or species extinctions^37^. The perceived importance of microbiome stability largely stems from studies of human and other organisms’ health where a dysbiotic, or unstable microbiome is considered a disease indicator^13^. However, disruption is natural in any system, therefore, defining the properties that maintain function despite disturbances are key. Network tools have their limitations for inferring specific functions, but there is mounting evidence that network complexity is often linked to stability in real world systems^1^. As demonstrated here, complexity cannot be defined by any one property and some are better predictors of robustness than others.

From our assessment of 33 networks spanning microbiomes inhabiting a range of spatial scales, environments and habitats, universal principles of microbiome robustness have emerged. In particular, robust networks were characterized by the maintenance of taxa that are highly connected and central within their co-occurrence networks, especially interkingdom networks with relatively higher proportions of predicted negative interactions. The role of these highly connected and central taxa in maintaining network architecture has parallels to keystone species in food webs, where their extinction has drastic impacts on communities and their functions^38^. While it is difficult to predict from taxa-based co-occurrence networks what functions of the microbiome might be compromised by keystone species’ extinctions, it is clear from our results that the diversity and composition of both fungal and bacterial communities would change significantly. For example, across the whole watershed, removing <10% of the bacteria and fungi with the highest betweenness centrality values led to a loss of >40% of all nodes, and similar patterns were observed across networks at all spatial scales (Fig. 2). While taxa with high betweenness centrality have similar roles in maintaining network structure, their identities among networks were not the same despite significant overlap in microbial community composition across hosts, habitats and the watershed (Fig. S17). Therefore, the shared specific properties of these taxa that encourage robustness deserves further investigation. Interkingdom interactions generally increased robustness, but this may again be a product of node-based properties such as node betweenness centrality and node degree, which were always significantly higher in interkingdom networks than single domains. However, betweenness centrality may be a stronger determinant of stability than node degree alone as previous studies have shown that only these nodes act as bridges connecting other highly central nodes, and their removal decreases network function^39^. Higher interkingdom node degree and betweenness centrality may be owed to fungi acting as connectors between modules in multi-kingdom assemblages^40^, possibly through the provision of physical niche space for bacterial colonization and dispersal^41^, or via metabolites that bacteria may exploit in nutrient-limited environments^42^. Therefore, despite fungal networks alone being least robust, the presence of fungi led to overall increased network stability.

We found interkingdom networks followed by bacteria and then fungi, to consistently harbor more predicted negative interactions, as well as a strong positive relationship between robustness and the proportion of negative edges in a given network. Whereas positive interactions have the potential to catalyze the mutual downfall of coupled species^13^, the prevalence of predicted negative interactions among more robust microbiome networks may be due to competition, predator-prey interactions, parasites or pathogens diffusing the effects of disturbances^43^ while keeping populations of detrimental species in check. For example, food web models put forth by Gross et al.^10^ found predator diversity to be a stabilizing factor by keeping prey populations under control. A similar result was also found in empirical food web research, where low predator-prey ratios tended to stabilize soil food webs^44^. Certain lineages of bacteria achieve this by suppressing pathogenic fungi through competitive root colonization, antifungal metabolite synthesis, or other biocontrol activities^45^. Pathogenic microbes themselves may also stabilize communities by promoting selected taxa and limiting the colonization of other microbes^46^. In our networks, fungal nodes assigned to *Candida albicans* always formed negative edges with *Weissella*, a genus of lactic acid bacteria with known antifungal activities that specifically inhibits *C. albicans* biofilm formation^47^. *Weissella* spp. also suppress pathogenic bacteria such as those in the genus *Acinetobacter*, and this negative link was also present in our networks^48^.

We set out to assess not only the effect of targeted and untargeted (random) node removal on microbiome robustness, but also the relationship between robustness and various measures of complexity. We use a definition of complexity, node richness and their edges, that parallels the ecological definition of species diversity and their interactions^1^. We find that while node (taxa) richness alone has a positive relationship with robustness, other additional measures of complexity are equally, if not more important for predicting stable microbiomes. Specifically, two related indices, connectance or the proportion of realized predicted interactions relative to all possible ones, and observed edge to node ratio. Both properties have previously been shown to be important for the stability of food webs^6^, social networks and cells^37^, but here we find they are also strong predictors of microbiome stability, even when accounting for the effect of richness on these relationships. So, while much emphasis has been placed on the importance of biodiversity for maintaining function, we suggest that additional consideration of interaction type (positive or negative) and interaction frequency is warranted.

Despite their stabilizing effect on bacterial networks, fungal networks alone were universally the least robust and defined by their high modularity, many positive edges and low node degree. High modularity is a network property that has repeatedly been associated with stability, purportedly due to the inability of disturbances to radiate beyond individual modules^14,31^. However, in the case of our robustness analyses which measured the remaining size of the largest network component (module) after node removal relative to its starting size, rapid module collapse may be due to the extinction of specific keystone taxa connecting multiple network components. This suggests that fungi connect sub-networks and potentially facilitate connectivity and resource sharing to a greater extent than bacteria^40^, but this increased communicability may be conferred at the expense of network stability^49^. Fungal networks were also composed primarily of positive edges (≥95%; Fig. 3), which could potentially explain their low robustness and predicted vulnerability to extinction cascades. Although cooperative mechanisms may be beneficial to the fitness of individual hosts, positive interactions are thought to destabilize ecological networks as perturbations can spread more rapidly when species are tightly linked in positive feedback loops^13,50^. For this reason, the loss of any one fungal species causes a more rapid deterioration of the network.

We assessed the complexity *x* stability relationship for a wide range of microbiomes found across land, sea and stream and inhabiting hosts ranging from birds to bugs to plants. While much prior attention has been placed on the value of biodiversity, specifically species diversity, in maintaining stable communities and their functions^2^, we find that additional aspects of complexity such as the frequency and type of interactions among species are equally if not more important predictors of robustness. Also, networks inclusive of the least robust microorganism networks, in this case fungi, generally increased the overall stability of bacterial networks, indicating that interkingdom co-occurrences are another important and often overlooked component of complexity that can positively affect stability. While stability may promote long-term coexistence of species^18,19^, other examples of stability in nature include less-favorable ecosystem states such as biological invasions^20^ and gut microbiota dysbiosis following antibiotic treatment^21^. Therefore, in the context of microbiome engineering it is critical to consider the properties of the reference system, whether it be a healthy gut, a productive agricultural field or an ecosystem, that are important to emulate, which may, or may not include stable microbial communities or stable functions of the microbiome. Indeed, enhancing the ability of microbiomes to acclimate or adapt rather than just persist may be an equally important aspirational trait for microbiome engineering and one that is only recently beginning to receive attention^51^. Future experiments assessing these principles are encouraged as this watershed-wide model study system has now provided clear testable hypotheses for the fundamental building blocks of microbiome stability.

## Methods

### Sampling Description

Our model watershed in the Waimea Valley, Oʻahu, Hawaiʻi U.S.A contains a precipitation gradient rivaling that of entire continents (change of ∼3.5 m in precipitation from the headwaters to estuary), where in less than 12 km rainfall levels at the driest and wettest sites match those observed in the driest portion of the African savanna to the Hoh Rainforest, WA, the wettest place in the continental United States. This gradient corresponds with additional dramatic changes in temperature and elevation (Fig. S1a). Microbial diversity was sampled across the entire Waimea watershed, from seven paired stream and terrestrial plots (20 m diameter) and seven marine plots from near-shore sand flats and coral reefs of the bay (21 plots total). From each plot, 113 + 54.5 (SD) biological samples were collected from host organisms and environmental substrates (Fig S1b). Sampling was roughly balanced across plots by trophic hierarchy (primary producers, consumers, or environmental substrates; Fig. S1c), and sample type followed the Earth Microbiome Project ontology (EMPO), which delineates samples by host association, salinity, and substrate type^11^. Of the most granular of the EMPO categories (EMPO3) we sampled 13 out of the 17 total. Thus, our sampling effort covers >75% of earth’s microbial habitats. Fungal and bacterial amplicons (ITS and 16S) were sequenced on an Illumina HiSeq run with 2 x 250 paired-end sequencing (Illumina Inc., San Diego, CA, USA). For full details on sampling, see^24^.

### Bioinformatics and statistics

Bacterial and fungal sequences were processed, filtered, and annotated using the Metaflow|mics pipelines^52^ as in ^24^. Sequences were clustered into 97% operational taxonomic units (OTUs) using the *uclust* function in QIIME version 1.9.1^53^. Here, we use 97% sequence similarity OTUs for both ITS and 16S, which represents species equivalents in the former^54^ and likely a slighter higher level of biological organization the latter^55^. As with any distance thresholds used for OTU construction, there will always be a tradeoff in terms of grouping or splitting sequencing reads^56^. Samples and OTUs with low abundance were removed, with cutoffs determined by “breaks” in distributions of log-transformed read counts by sample and OTU. For fungi, this entailed culling samples with 190 or fewer reads and OTUs with 4 or fewer reads. For bacteria, we culled samples with 3,000 or fewer reads and OTUs with 5 or fewer reads. Because OTUs with low prevalence can lead to the formation of spurious edges in co-occurrence networks^31^ we filtered out OTUs present in fewer than 20% of samples in the whole dataset within a given sample type (EMPO3 designation). We also removed OTUs present in only one sample, as these OTUs could not co-occur with other OTUs. The sums of prevalence-filtered OTUs were kept in a separate row to maintain overall sample counts in network inference, and this row was removed for downstream network visualization and analysis. Each resulting dataset consisted of 1,384 samples with 2,128 OTUs in the ITS dataset, and 13,468 OTUs in the bacterial 16S dataset. Spatial autocorrelation was assessed for each locus (16S and ITS) using a Mantel test on a Bray-Curtis distance matrix of OTU abundances and a geographic distance matrix. Significance was determined with the Spearman correlation coefficient and 999 permutations. The overlap of OTUs among EMPO3 sample types for bacteria and fungi was visualized using venn diagrams (Fig. S8).

### Network Construction

All analyses were conducted in R v.4.0.0. Co-occurrence networks were constructed using the *SpiecEasi* package^57^, which considers the compositional nature of microbial data, is suitable for datasets in which OTUs outnumber samples, and is robust against false positives, outperforming methods such as traditional Pearson correlations or *SparCC*. *SpiecEasi* was also chosen for its ability to handle interkingdom data by applying the center-log ratio transformation to each dataset before concatenation, which satisfies the assumptions of equations used to generate the inverse covariance matrix^58^. All networks (single- and interkingdom) were constructed using the Meinshausen and Bühlmann (“MB”) method on our prevalence-filtered abundance tables, with a pulsar parameter threshold of 0.01 and screening parameter set to TRUE to account for large OTU counts. A lambda minimum ratio of 1e−5 was used unless otherwise specified. Rather than specifying correlation thresholds for edge formation, *SpiecEasi* infers edges by conditional independence, where an edge can only exist between two nodes given all other nodes in the network. That is, if a relationship between two nodes can be explained by an external taxon, an edge will not be inferred, reducing the incidence of indirect edges^59^. Rather than with a false discovery rate, the fidelity of a network generated in *SpiecEasi* comes from the process of sparsity tuning^57^. A graph solution path from empty to complete is produced, and 80% subsamples of the data are randomly and repeatedly taken to estimate the full solution path. The final selected graph has the most stable edge incidences across subsamples, based on an optimized lambda value balancing sparsity and model fit, where sparser networks indicate less variable edges. Edge sign (positive or negative) is taken from the regression coefficients from *SpiecEasi*^57^.

A common concern with network interpretation is the unknown influence of abiotic factors on edge inference, in that it is possible for edges to form from common responses to environmental factors rather than actual species interactions^60^. One suggestion to mediate this issue is to hold these variables constant among constructed networks^60^. However, this leads to context dependency thereby limiting the inference of universal microbiome properties (i.e., one could say that a property is stabilizing in high-saline environments, but not all environments). Because our goal was to identify fundamental attributes of microbiome stability regardless of environmental influence we deemed “universal” properties those that were consistent across networks representing (1) the whole watershed (2) its intrinsic diversity of environments as well as (3) sample types among these environments and between microbial kingdoms.

Networks representing the entire watershed were constructed from the full fungal and bacterial datasets with a lambda minimum ratio of 1e-2 to account for large OTU counts. Resulting full fungal, bacterial, and interkingdom networks were represented by 2,128 OTUs, 13,468 OTUs, and 15,596 OTUs, respectively. The full datasets were also used to construct networks representing individual habitats and sites along the terrestrial gradient, but were subject to separate culling measures to address differences in sequencing depth before network assembly. From each habitat (marine, stream, and terrestrial), we randomly selected the lowest common number of samples (17) across five distinct EMPO3 categories kept roughly standardized by trophic guild, for a total of 85 samples per network. From each terrestrial gradient site (Beach, Estuary, Entrance, Confluence, Waterfall, Drum Road, and Ridge), we randomly selected the lowest common number of samples within an EMPO3 category across all sites, for a total of 46 samples per network. The resulting OTU counts for each habitat and gradient network are listed in Table S1, and generally range from 918 to 9,061 OTUs in habitat networks, and 721 to 6,627 OTUs in gradient networks. To assess the influence of richness on observed network properties, trimmed networks for each scale (watershed, habitat, and terrestrial gradient) were also generated by culling networks to the lowest common number of nodes (721 nodes).

### Network Characterization

Networks were analyzed using the *igraph* package^61^. To assess whether the inferred associations differed from random expectations (e.g., Poisson node degree distribution), we tested each network for small-world and scale-free patterns^39,62^. Node degree (*k*, the number of connections to a node) was calculated for each network and the distribution of node degree (*P(k)*) was used for the assessment of whether networks are scale-free. Node betweenness centrality (*g*) or how often a node occurs along the shortest path between other nodes was used for the assessment of node contribution to network robustness. Complexity as measured by node count (richness) and edge to node ratio, along with modularity (the degree to which a network partitions into distinct and highly connected sub-communities) and connectance (*C*, the number of realized edges in a network among all the possible ones), were also calculated. The relationship between node degree and betweenness centrality for all networks was examined via linear regression for all networks (whole watershed, habitat and gradient; Fig S9) and significant differences in node-level metrics (node degree and betweenness centrality) were calculated with ANOVA and Tukey’s honestly significant difference tests. Modules were identified using the *rnetcarto* package^63^, and we calculated percentages of EMPO3 categories per module by taking the samples associated with a given module’s OTUs and the EMPO3 categories from which they came. Negative edges were identified using the beta matrix from *SpiecEasi*^57^, and the percentage of negative edges was calculated for each network.

### Robustness Analyses

Extinction cascades or food web collapses are often linked to the loss of specific species with a relatively large number of interactions with other species^64^. Similarly, we can quantify microbiome network stability via attack robustness in which nodes (microbial taxa) are sequentially removed in order of their relative betweenness centrality or at random. Then, we can measure the size of the largest remaining connected component in the network and divide this value by the starting size of the largest component as an indicator of network stability based on the remaining ability of nodes to interact with one another^28,39^. Robust networks are considered those with connected structures maintained despite the loss of nodes, which is indicated by a larger area under the curve (AUC)^58^.

Robustness curves were calculated using the *brainGraph* package^65^. We generated three curves for each network: random node knockout (error attack), and two forms of targeted attack: nodes with the highest or lowest betweenness centrality, calculated iteratively after each knockout^37^. Robustness curves were generated for all single- and interkingdom networks (whole-watershed, habitat, and the terrestrial gradient) and compared within network type. To examine the relationships between robustness and complexity (as measured by richness, the ratio of edges to nodes, connectance, modularity, and percent negative edges) we plotted the AUC for each fungal, bacterial, and interkingdom networks for each scale (watershed, habitat, and gradient), and performed linear regression.

### Phylogenetic Diversity of Bacterial Communities

To examine the relationship between bacterial phylogenetic diversity (a form of complexity) and robustness, phylogenetic diversity was calculated as the mean pairwise phylogenetic distances (MPD) between OTUs within every sample present in a bacterial network (i.e., MPD values were not derived from networks themselves). This analysis was not done for fungal samples because the ITS locus is less phylogenetically informative. Standardized effect sizes (SES) of phylogenetic community structure were computed by comparing observed MPD values to MPD values expected under a null model where the taxa labels of each sample’s distance matrix were randomized, and iterated 999 times. Calculating SES values, as opposed to MPD alone, allows us to examine whether co-occurring OTUs are more or less related than expected by chance, across habitats and sites along the terrestrial gradient. Negative SES values indicate greater phylogenetic clustering, while positive values indicate phylogenetic dispersion^66^. Calculations were done using the *picante* package^66^, with the original distance matrix computed using cophenetic in base R. Significant differences in MPD between habitats or sites along the terrestrial gradient were determined with ANOVA and Tukey’s HSD tests.

## Data Availability

DNA sequences and project metadata are archived in the NCBI Sequence Read Archive (SRA) underBioProject accession no. PRJNA701450 and in Qiita under study ID 13115.

## Code Availability

All data and code for analyses are located at https://github.com/kkajihara/waimea_stability.

## Acknowledgements

Many thanks to Eoin Brodie, Pieter Dorrestein, Jannet Janssen, Rob Knight, Jennifer Martiny, Monique Chyba, and Edward Ruby for their input; Cedric Aridakessian for assistance with data processing; Laura Tipton for lending her network expertise; and Joshua Buchanan, Kahiwahiwa Davis, Brennan Hee, Tanja Lantz Hirvonen, Reece Kilbey, Terrance McDermott, Joma Santos, Leina Uemura, Nicole Yoneishi, Anastasia Morse, Shayle Matsuda, Campbell Gunnel, David Pence, Chris Wall, and Jeff Kuwabara for assistance in the laboratory and field. We also thank Richard Pezzulo, Chad Durkin, Josie Hoh, and Laurent Pool of Hiʻipaka LLC and Waimea Botanical Garden for their assistance, and the Hauʻoli Mau Loa Foundation for fellowship support to KTK. The technical support and advanced computing resources from University of Hawaii Information Technology Services – Cyberinfrastructure, funded in part by the National Science Foundation CC* awards 2201428 and 2232862 are gratefully acknowledged. Funding for this work was provided by the W.M. Keck Foundation, the office of the Vice Chancellor for Research at the University of Hawaiʻi at Mānoa to C-MĀIKI (Center for Microbiome Analysis through Island Knowledge and Investigation, the University of Hawaiʻi Sea Grant, NIH award P20GM125508, and NSF awards 2124922 and 2023298.

## Contributions

KTK and NAH conceptualized the idea, supervised the project, collected and processed samples, collected and analyzed the data and wrote the manuscript; MY analyzed the data and reviewed and edited the manuscript; ASA supervised the project and reviewed and edited the manuscript; NC collected samples; JLD analyzed the data and reviewed and edited the manuscript; KMSF collected samples; KLF supervised the project, collected samples; MM-N supervised the project and reviewed and edited the manuscript; MM supervised the project, collected samples and reviewed and edited the manuscript; KKN supervised the project, collected samples and processed them; CEN supervised the project, collected samples, collected the data and analyzed it, reviewed and edited the manuscript; RLR collected and processed samples; WJS collected and processed samples and reviewed and edited the manuscript; SOIS supervised the project, collected samples and processed them and collected the data; MAT collected samples; JYY supervised the project and collected the data; DY processed samples and collected the data.

The authors declare no competing financial interests.

