## Supplementary Information for "Diversity, connectivity and negative interactions define robust microbiome networks across land, stream, and sea"

### I. Supplementary Figures

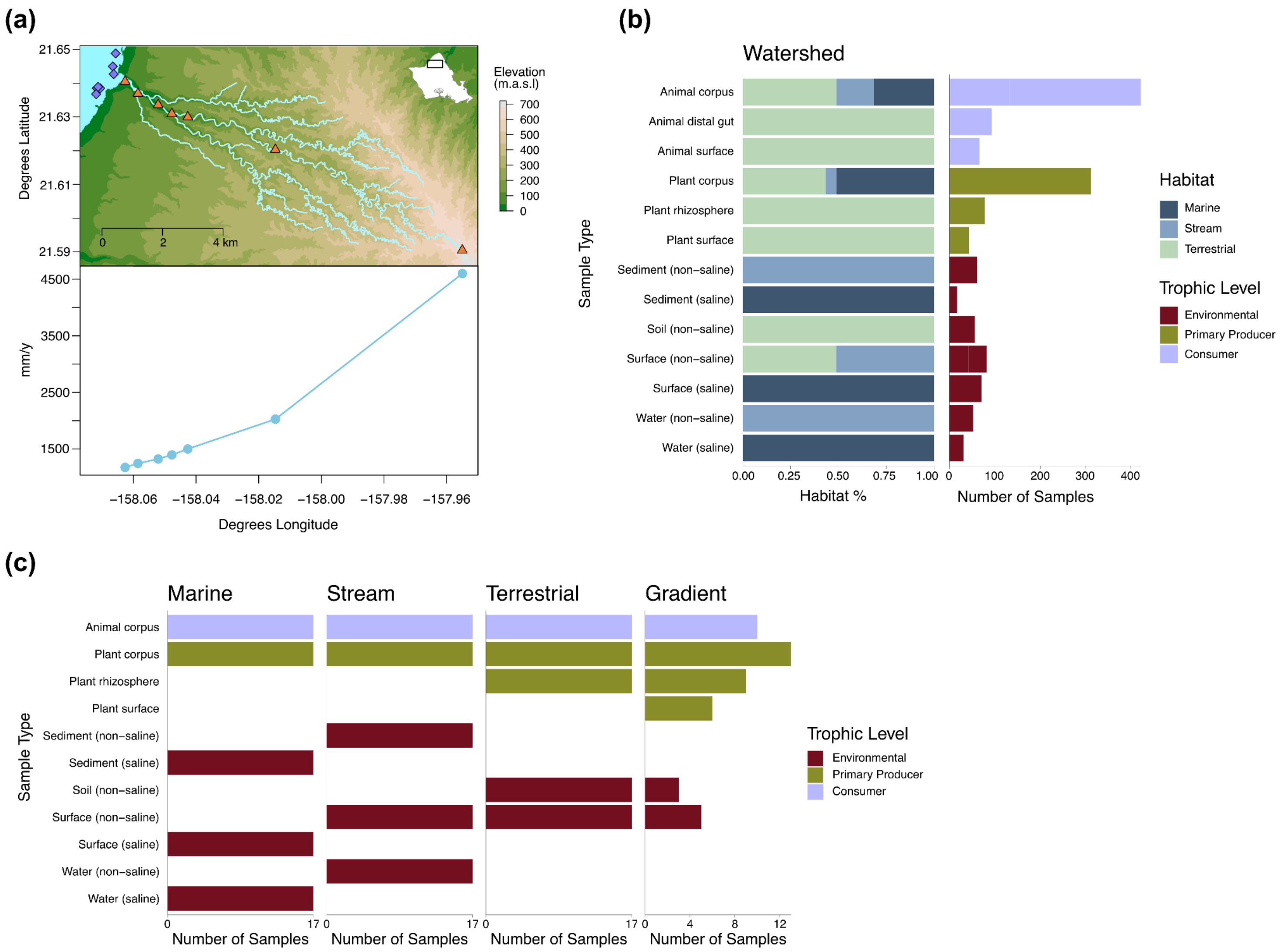

Figure S1. Site locations and sampling information by network type, adapted from[1]. (a) Positions of plots (n = 21) within the Waimea watershed (O‘ahu, Hawai‘i). Blue diamonds are marine habitat plots, and orange triangles are paired stream/terrestrial habitat plots. The elevation gradient is represented in the top panel (“m.a.s.l” indicates meters above sea level) and the rainfall gradient in the bottom panel in millimeters per year. (b) Habitat and trophic level associations of samples included in the watershed-level networks (n = 1,384). Sample types are described by the Earth Microbiome Project Ontology (EMPO Level 3). (c) Trophic level associations of samples included in the habitat networks (marine, stream, and terrestrial; n = 85 for each habitat), and the terrestrial gradient network (far right; n = 46 for each site along the gradient). Note changes in values on the x-axes.

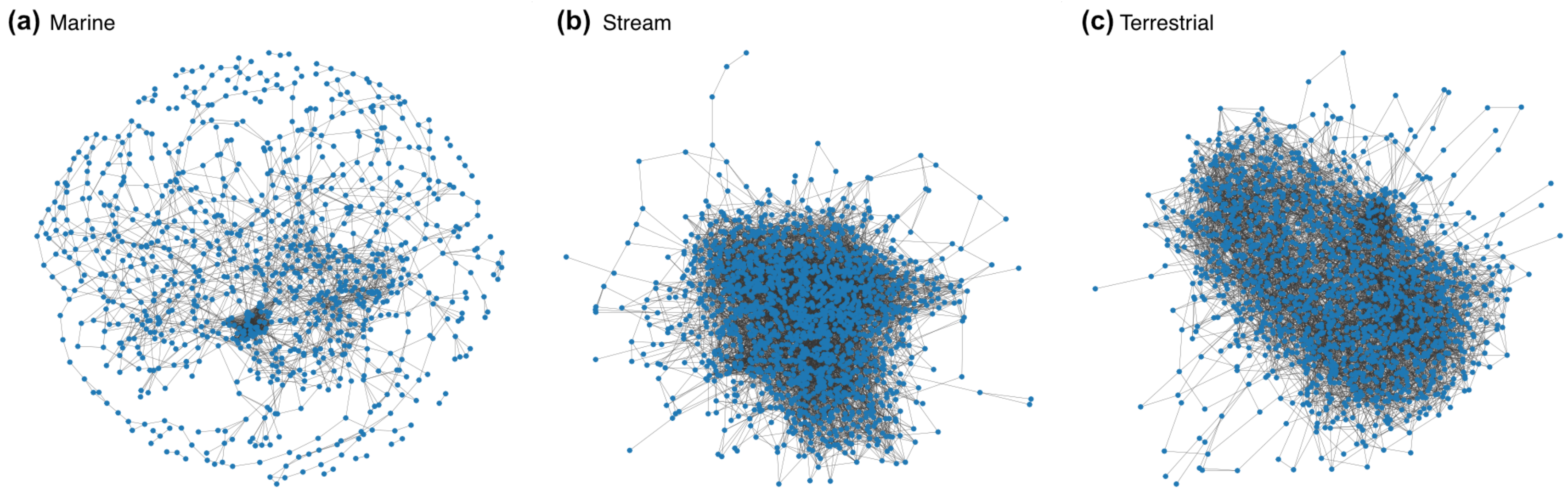


Figure S2. Fungal network visualizations from each habitat in the watershed.

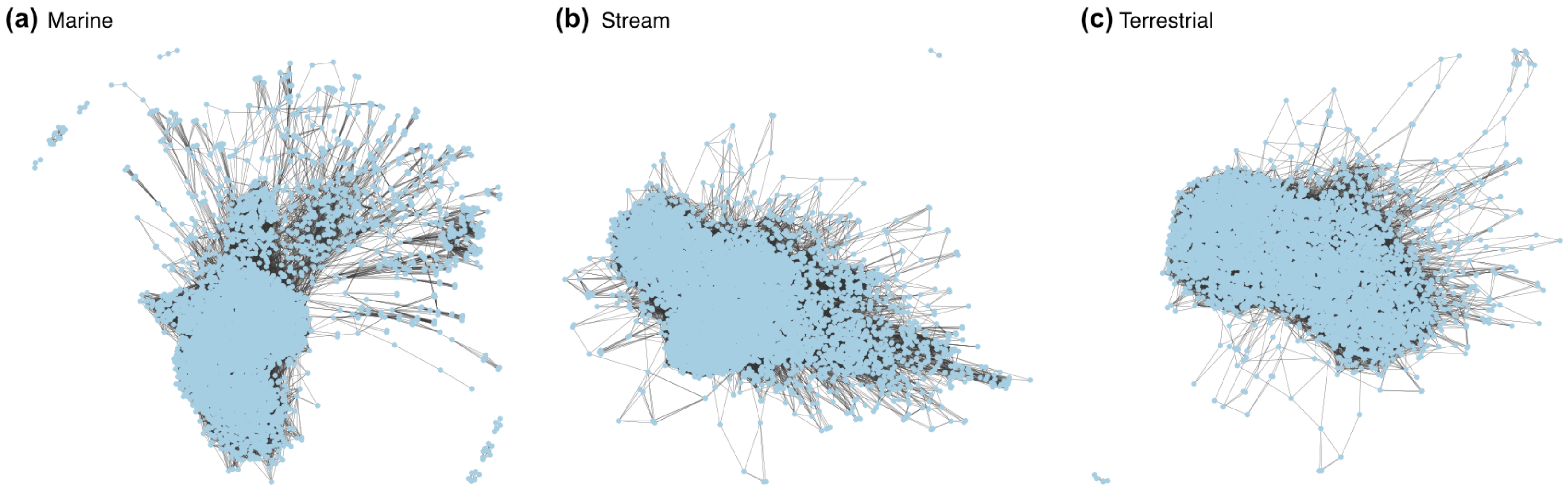


Figure S3. Bacterial network visualizations from each habitat in the watershed.

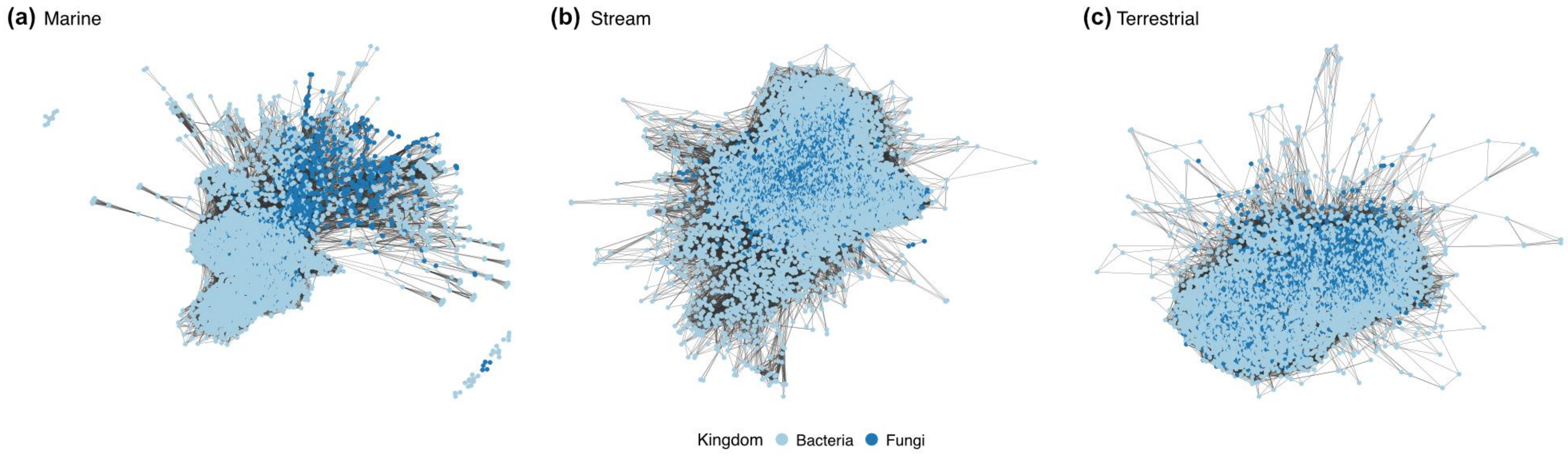


Figure S4. Interkingdom network visualizations from each habitat in the watershed, light blue circles represent bacterial taxa and dark blue fungi.

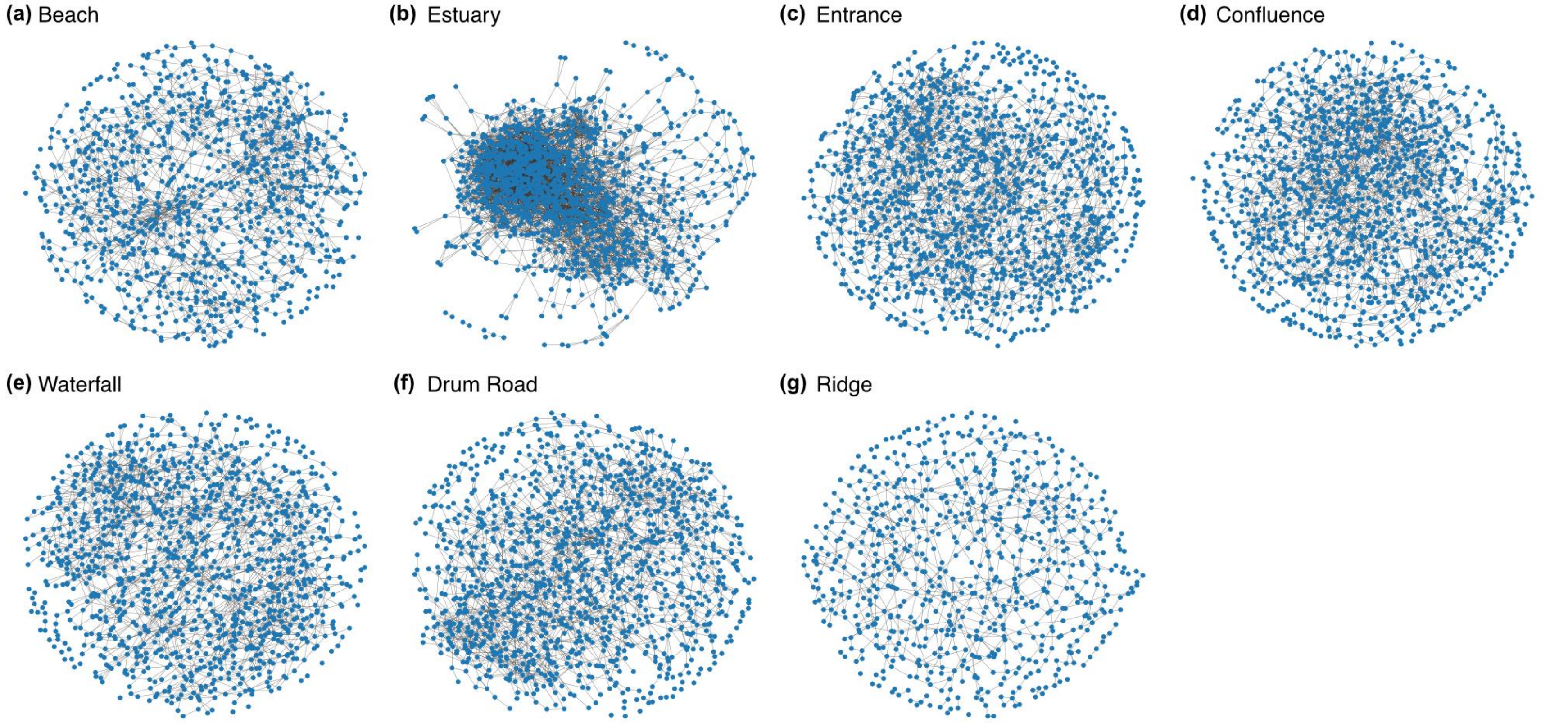


Figure S5. Fungal network visualizations from each site along the terrestrial habitat gradient, in order from the mouth to the headwaters of the watershed (a-g).

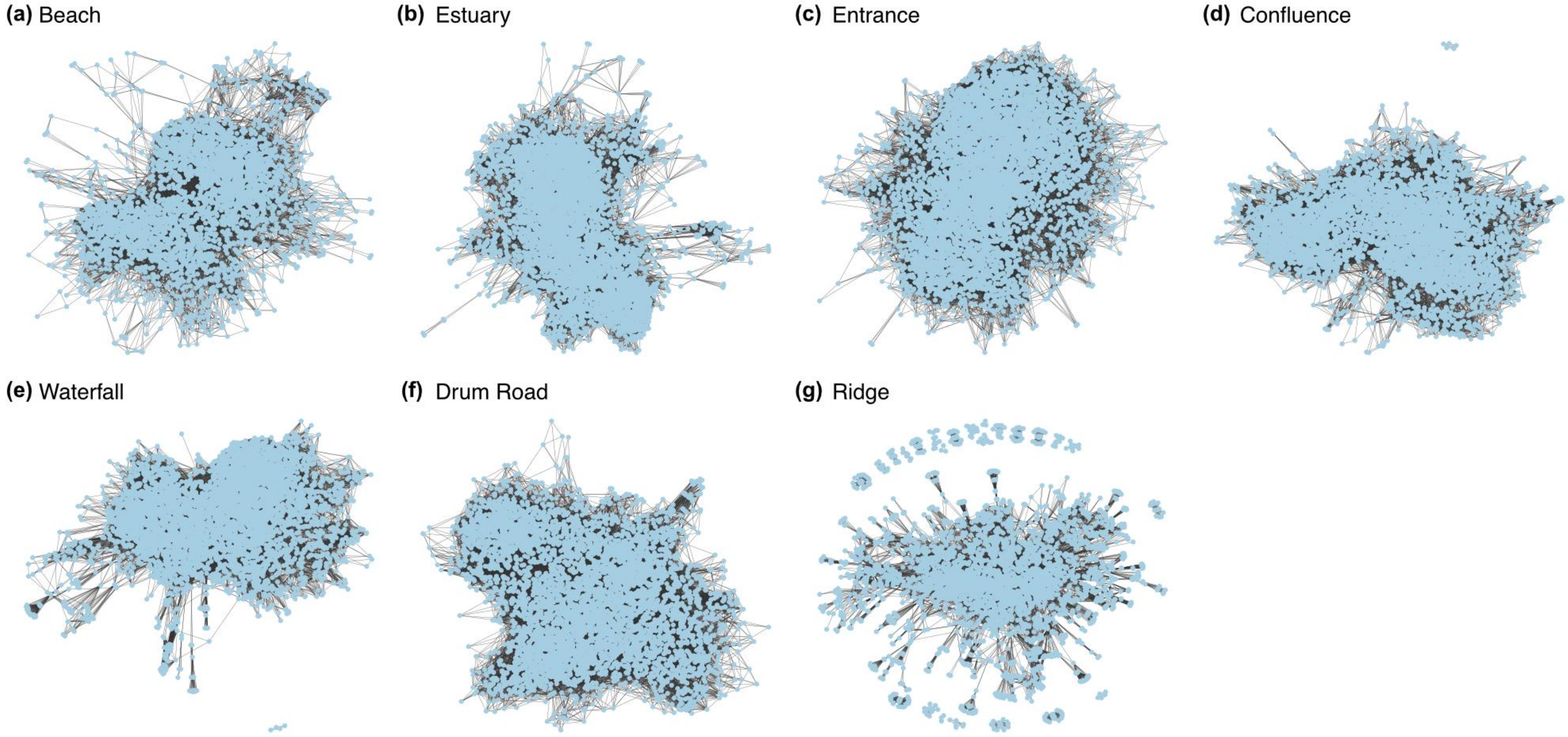


Figure S6. Bacterial network visualizations from each site along the terrestrial habitat gradient, in order from the mouth to the headwaters of the watershed (a-g).

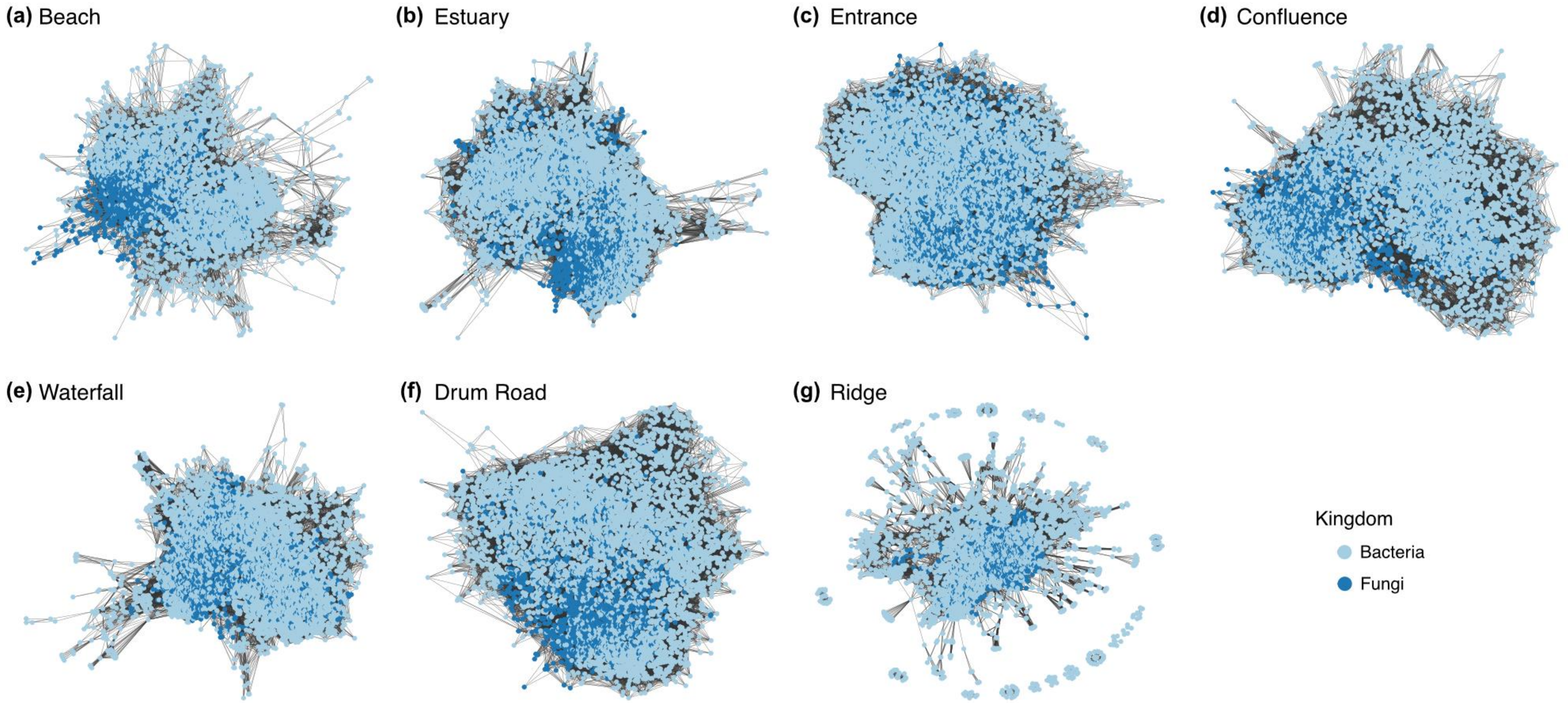


Figure S7. Interkingdom network visualizations from each site along the terrestrial habitat gradient, in order from the mouth to the headwaters of the watershed (a-g). Light blue circles represent bacterial taxa and dark blue fungi.

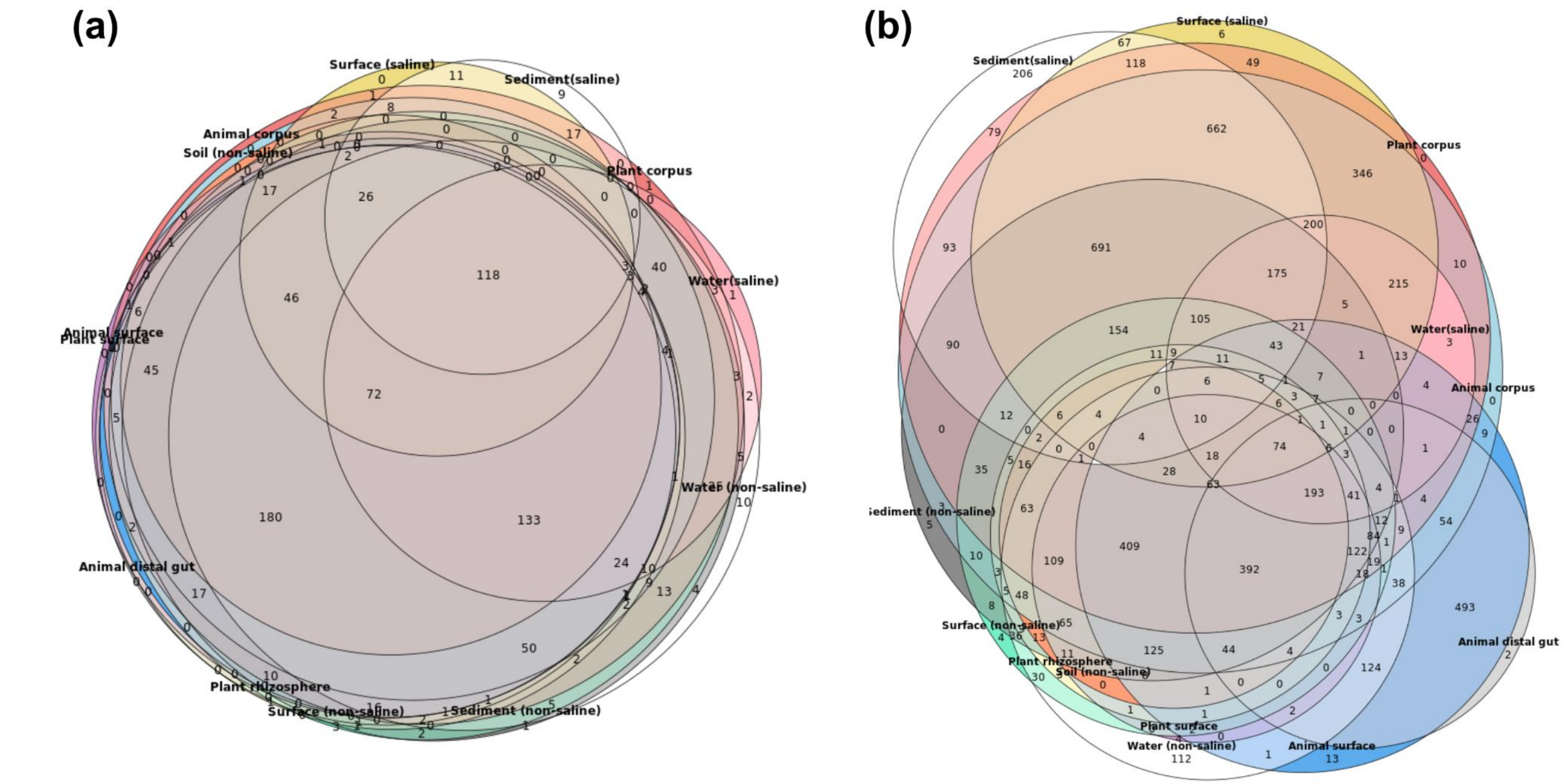


Figure S8. Operational Taxonomic Unit (OTU) identity by sample type (Earth Microbiome Project Ontology Level 3) for fungi (a) and bacteria (b) from the entire Waimea Watershed consisting of 1,384 samples with 2,128 OTUs in the fungal ITS dataset, and 13,468 OTUs in the bacterial 16S dataset.

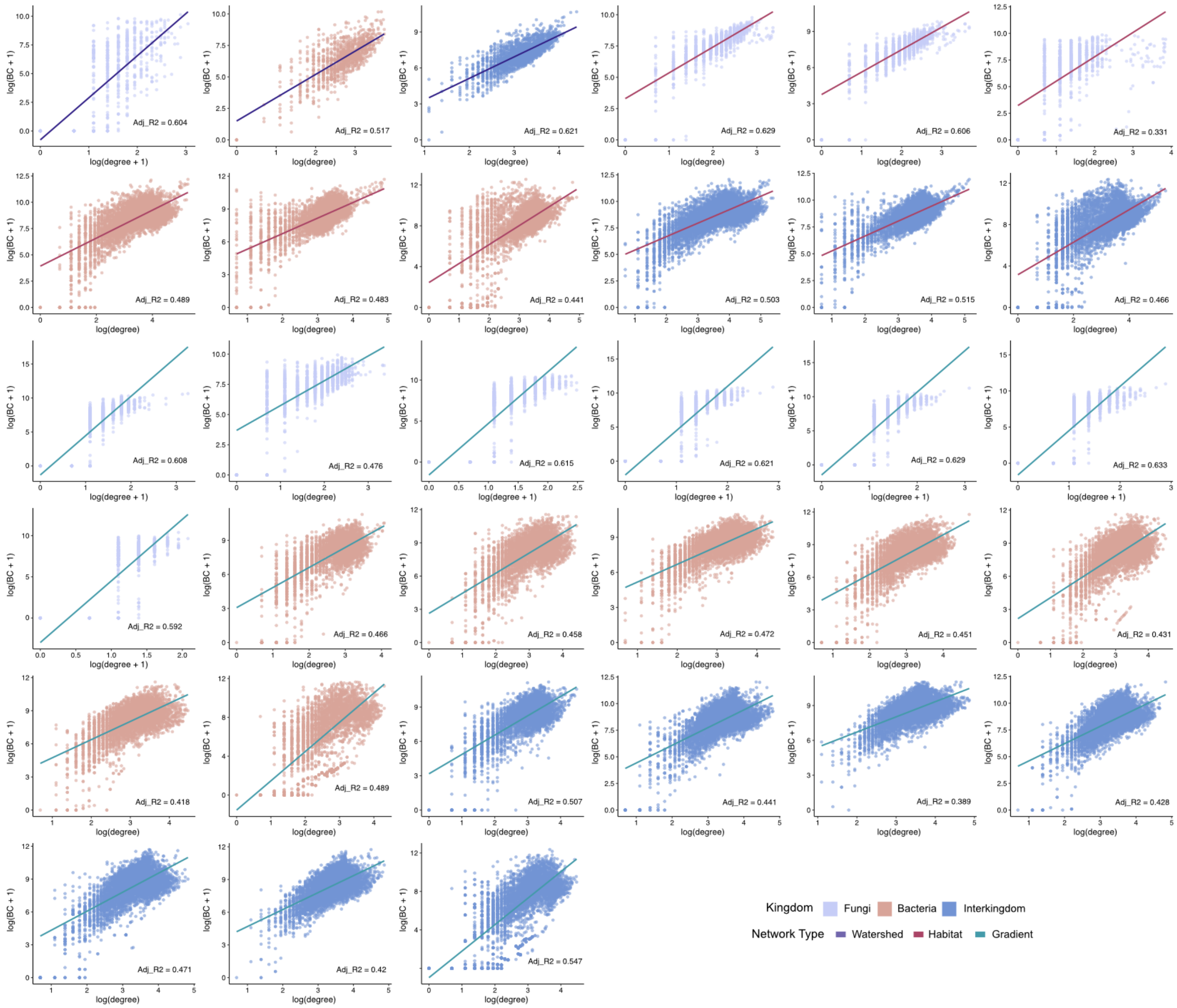


Figure S9. Relationship between node degree and betweenness centrality assessed via linear regression for each of 33 total networks. Color of the dots represent either fungal (lavender), bacterial (pink) or interkingdom (blue) networks. Regression lines represent statistically significant relationships between node degree and betweenness centrality and their color represents spatial scale ranging from the whole watershed (purple), to habitats inclusive of marine, stream and terrestrial (red) or sites along a steep environmental gradient within the watershed (green) at an $\alpha \leq 0.05$ and in all cases $p < 0.001$.

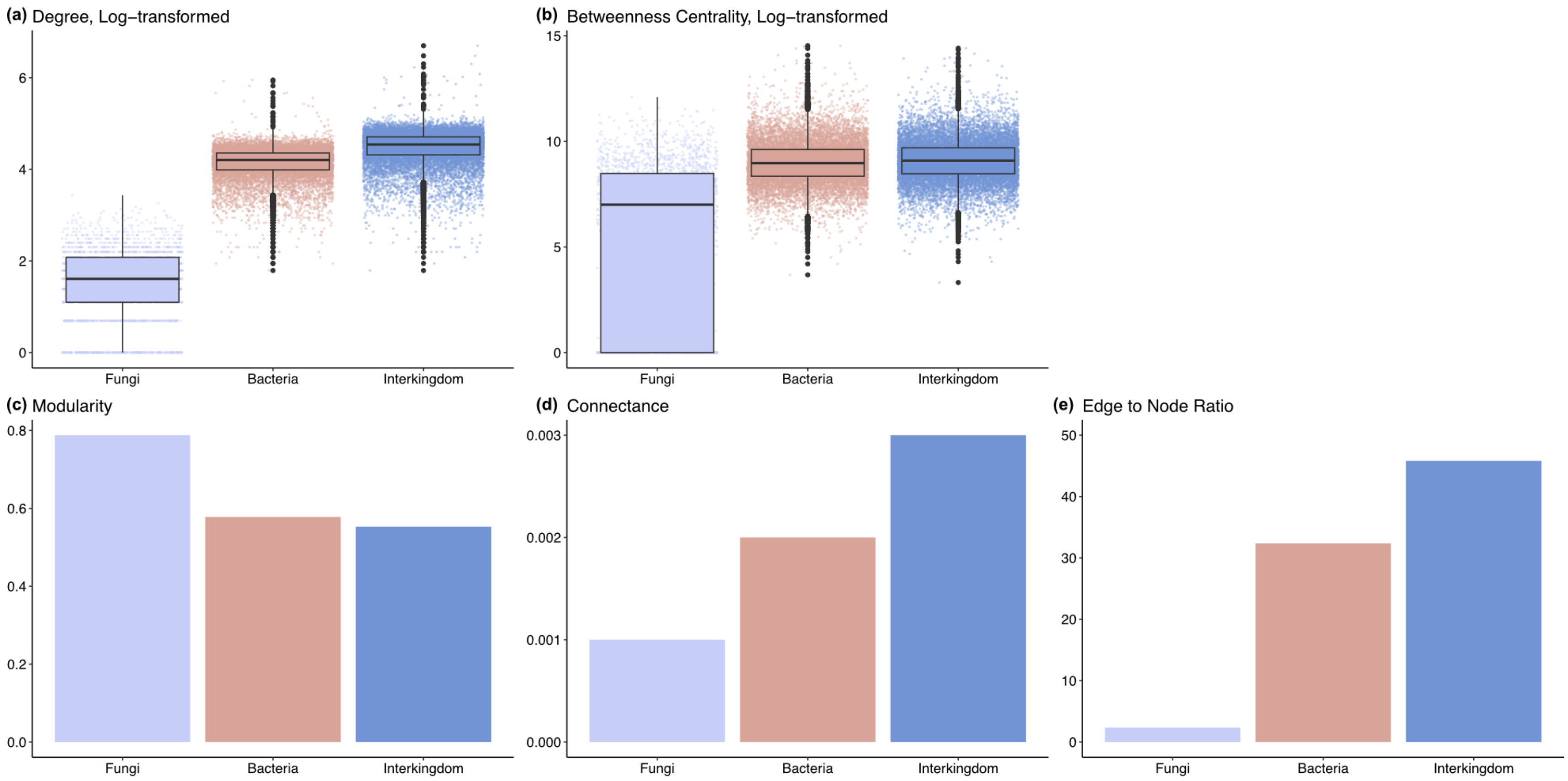


Figure S10. Network topologies representing complexity of the watershed networks at the node level (a-b) and whole network level (c-e). For panels a and b the center line within each box represents the median, whiskers represent the lower and upper 1.5 interquartile range, and black dots are outliers.

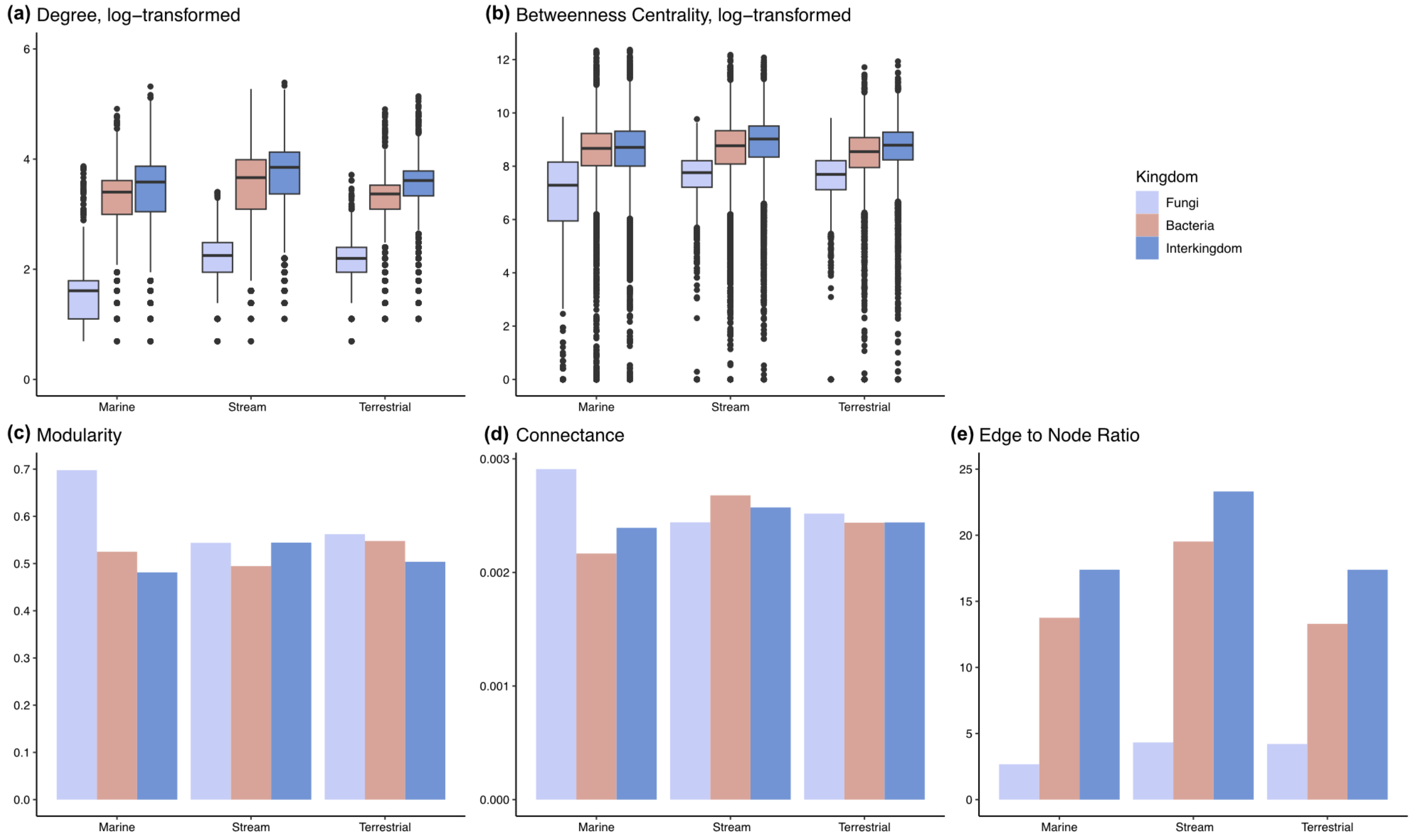


Figure S11. Network topologies representing complexity of the habitat networks at the node level (a-b) and whole network level (c-e). For panels a and b the center line within each box represents the median, whiskers represent the lower and upper 1.5 interquartile range, and black dots are outliers.

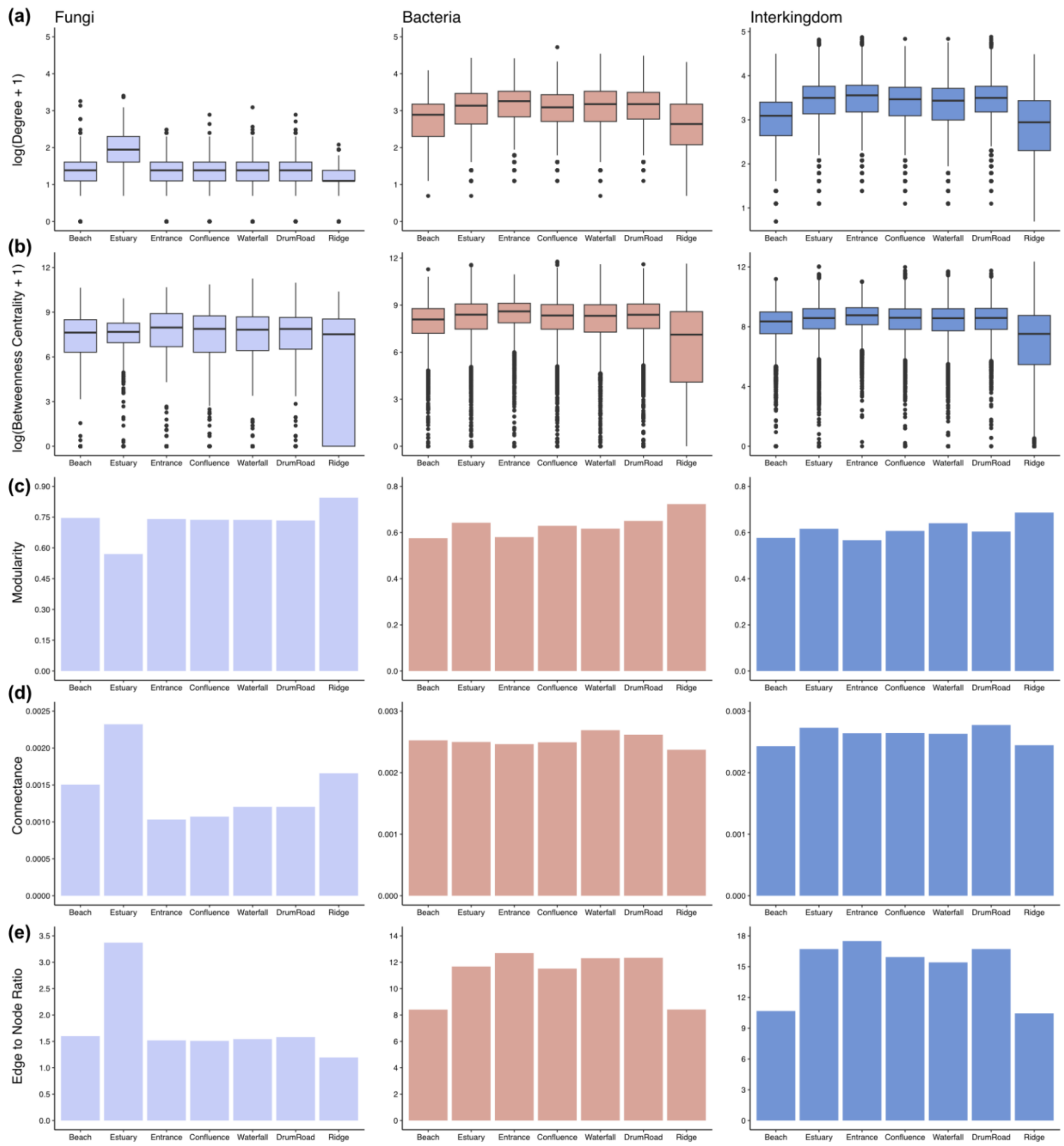


Figure S12. Network topologies representing complexity of the gradient site networks at the node level (a-b) and network level (c-f) in fungi (left column), bacteria (middle column), and interkingdom (right column) networks. Bars represent sites along the terrestrial gradient, in order from the mouth to the headwaters of the watershed. For panels a and b the center line within each box represents the median, whiskers represent the lower and upper 1.5 interquartile range, and black dots are outliers.

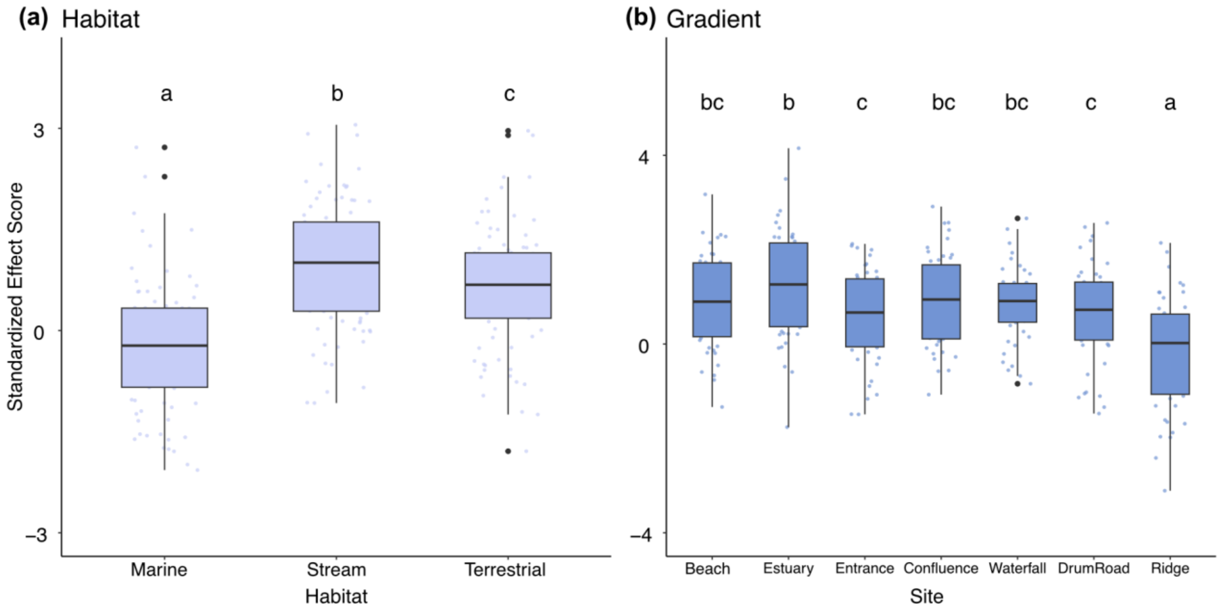


Figure S13. Phylogenetic distances among bacteria based on 16S amplicon sequencing from three habitats (a; marine, stream and terrestrial) and across a strong environmental gradient from the mouth of a watershed to its headwaters (b). Phylogenetic diversity was calculated as the mean pairwise phylogenetic distances (MPD) between Operational Taxonomic Units (OTUs) within every sample present in a bacterial network. Standardized effect sizes (SES) of phylogenetic community structure were computed by comparing observed MPD values to MPD values expected under a null model where the taxa labels of each sample's distance matrix were randomized, and iterated 999 times. Calculating SES values, as opposed to MPD alone, allows us to examine whether co-occurring OTUs are more or less related than expected by chance, across habitats and sites along the terrestrial gradient. Negative SES values indicate greater phylogenetic clustering, while positive values indicate phylogenetic dispersion. The center line within each box represents the median, whiskers represent the lower and upper 1.5 interquartile range, and black dots are outliers. Letters represent statistically significantly different habitats or sites (see Tables S9 and S10).

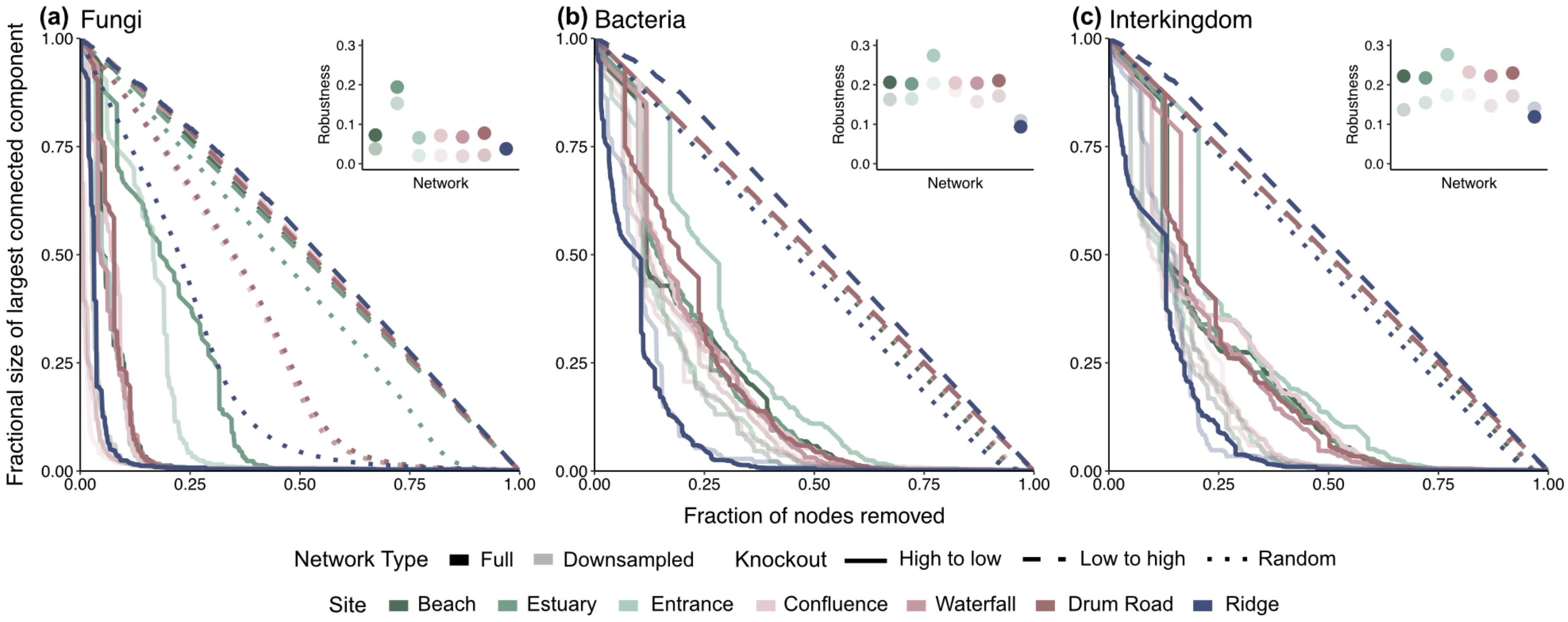


Fig S14. Attack robustness of microbial co-occurrence networks representing each site along a steep environmental gradient within an entire watershed for fungi (a), bacteria (b) or interkingdom networks (c). Robustness is measured as the size of the largest remaining network component relative to its starting size (which in this case included an average of 97.82% SD±3.00% of all nodes) after nodes are removed in order of high betweenness centrality (dark solid lines), low betweenness centrality (dashed lines), or at random (dotted lines). The lightened lines on each panel represent removal of nodes with high betweenness centrality from downsampled networks each with the same number of nodes as the smallest network (721). Each line represents a network from a site ranging from the mouth to the headwaters of the watershed. More robust networks are indicated by a larger area under the curve. The dots in each subpanel represent each networks' robustness metric as measured by area under the curve.

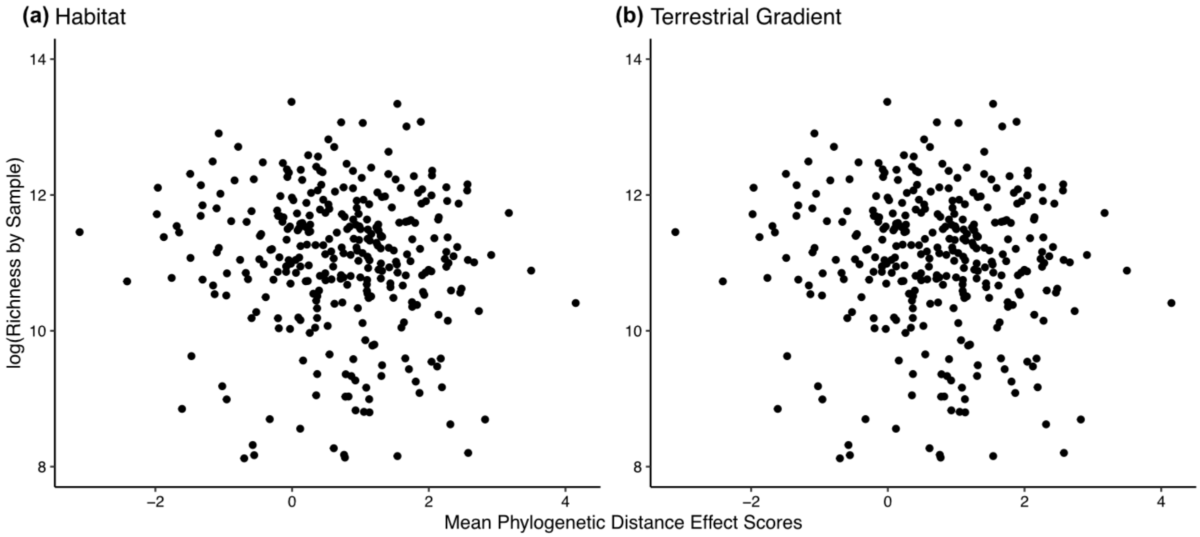


Figure S15. Scatter plots of bacterial operational taxonomic unit (OTU) richness versus mean phylogenetic distances across (a) habitats ($R^2$ = 0.006, $P$ = 0.217) and (b) sites along the terrestrial habitat gradient ($R^2$ = 0.004, $P$ = 0.283).

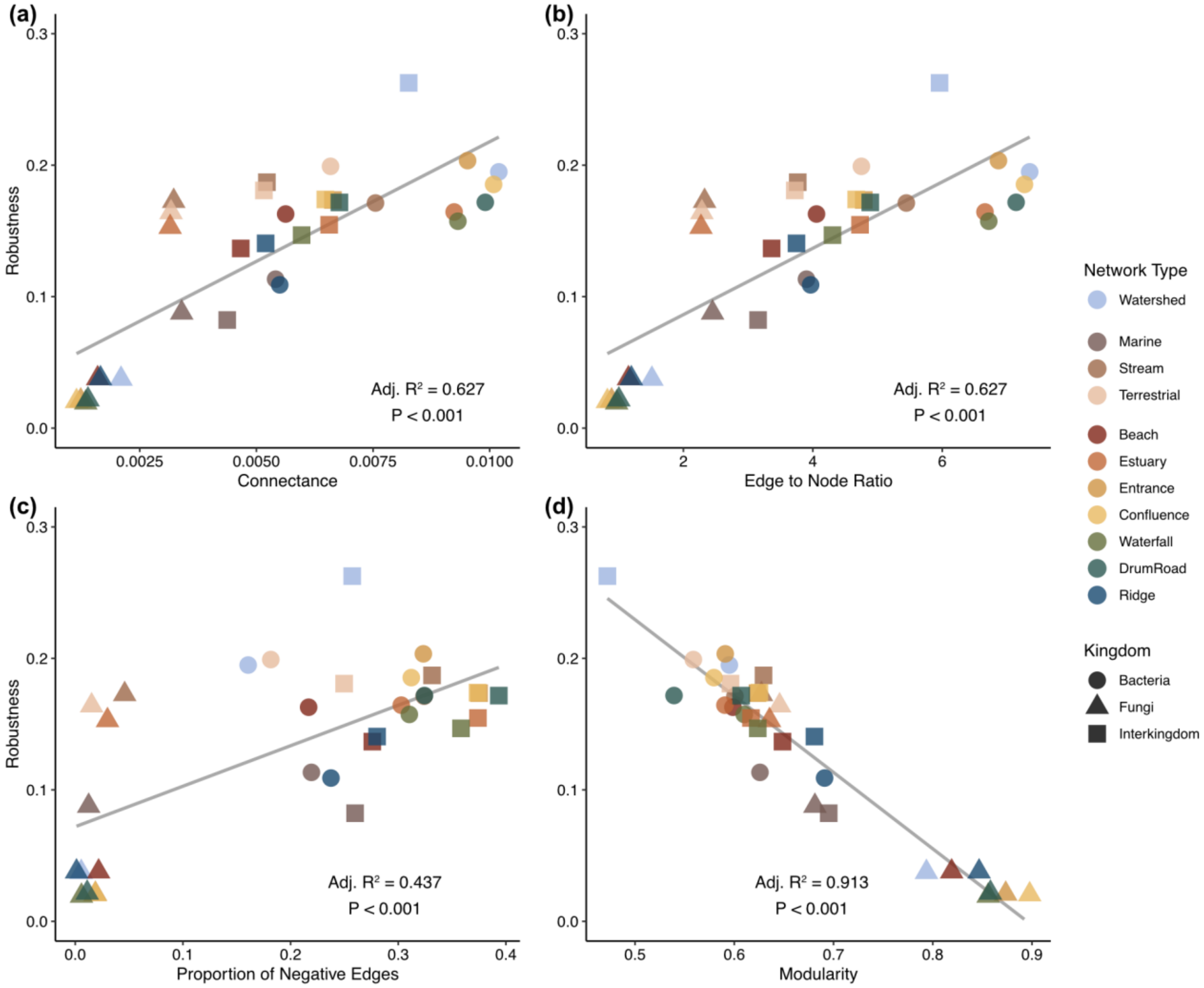


Figure S16. Linear regressions examining the relationship between robustness (area under the curve from figure 2) and various measures of complexity when node count (richness) is held constant to the smallest network size of 721 nodes. Each symbol represents either bacterial (circle), fungal (triangle) or interkingdom networks (squares) and each color a network of different spatial scale. All relationships are statistically significant at $\alpha \leq 0.05$ and $p<0.001$ to $p=0.001$.

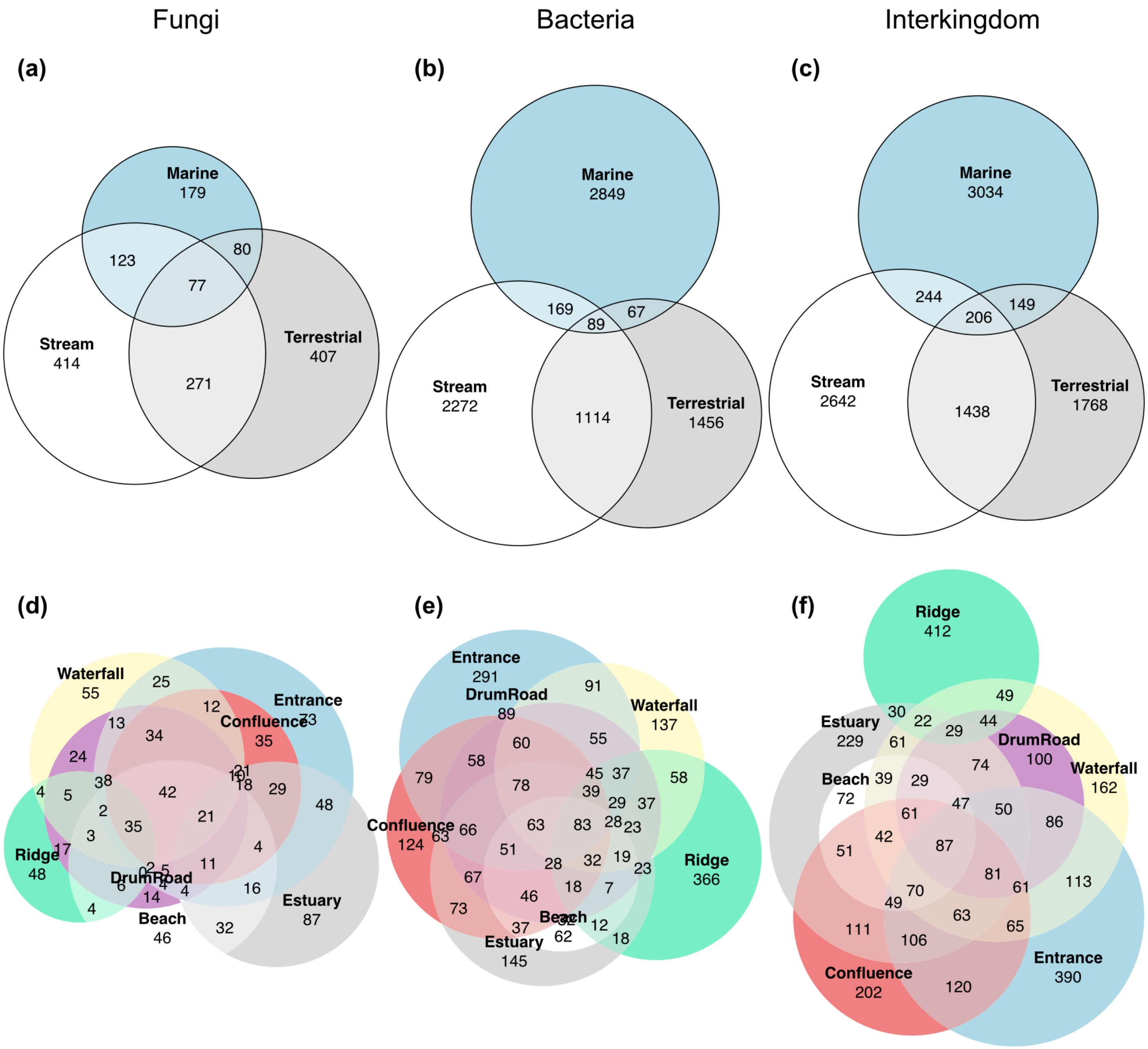


S17. Venn diagrams representing the overlap of the top 50% of nodes from each habitat (a-c) and each gradient site (d-f) with highest betweenness centrality values for fungi, bacteria and interkingdom networks. While there are a subset of nodes (taxa) with high betweenness centrality shared across habitats and sites, many are unique to their particular network.

#### II. Supplementary Tables

Table S1. Counts of operational taxonomic units (OTUs) by network.

| Type | Kingdom | Habitat | Site | Samples | OTU Richness |
|---|---|---|---|---|---|
| Watershed | Fungi | All | All | 1384 | 2,128 |
| Watershed | Bacteria | All | All | 1384 | 13,468 |
| Watershed | Interkingdom | All | All | 1384 | 15,596 |
| Habitat | Fungi | Marine | All | 85 | 918 |
| Habitat | Bacteria | Marine | All | 85 | 6349 |
| Habitat | Interkingdom | Marine | All | 85 | 7267 |
| Habitat | Fungi | Stream | All | 85 | 1772 |
| Habitat | Bacteria | Stream | All | 85 | 7289 |
| Habitat | Interkingdom | Stream | All | 85 | 9061 |
| Habitat | Fungi | Terrestrial | All | 85 | 1671 |
| Habitat | Bacteria | Terrestrial | All | 85 | 5453 |
| Habitat | Interkingdom | Terrestrial | All | 85 | 7124 |
| Gradient | Fungi | Terrestrial | Beach | 46 | 1063 |
| Gradient | Bacteria | Terrestrial | Beach | 46 | 3330 |
| Gradient | Interkingdom | Terrestrial | Beach | 46 | 4393 |
| Gradient | Fungi | Terrestrial | Estuary | 46 | 1452 |
| Gradient | Bacteria | Terrestrial | Estuary | 46 | 4672 |
| Gradient | Interkingdom | Terrestrial | Estuary | 46 | 6124 |
| Gradient | Fungi | Terrestrial | Entrance | 46 | 1472 |
| Gradient | Bacteria | Terrestrial | Entrance | 46 | 5155 |
| Gradient | Interkingdom | Terrestrial | Entrance | 46 | 6627 |
| Gradient | Fungi | Terrestrial | Confluence | 46 | 1410 |
| Gradient | Bacteria | Terrestrial | Confluence | 46 | 4617 |
| Gradient | Interkingdom | Terrestrial | Confluence | 46 | 6027 |
| Gradient | Fungi | Terrestrial | Waterfall | 46 | 1283 |
| Gradient | Bacteria | Terrestrial | Waterfall | 46 | 4575 |
| Gradient | Interkingdom | Terrestrial | Waterfall | 46 | 5858 |

| | | | | | |
|---|---|---|---|---|---|
| Gradient | Fungi | Terrestrial | DrumRoad | 46 | 1313 |
| Gradient | Bacteria | Terrestrial | DrumRoad | 46 | 4713 |
| Gradient | Interkingdom | Terrestrial | DrumRoad | 46 | 6026 |
| Gradient | Fungi | Terrestrial | Ridge | 46 | 721 |
| Gradient | Bacteria | Terrestrial | Ridge | 46 | 3549 |
| Gradient | Interkingdom | Terrestrial | Ridge | 46 | 4270 |

Table S2. Degree distribution tests of non-randomness for networks that did not have power law degree distributions. Power law fits of p > 0.05 (KS test) indicate that networks are scale-free (non-scale-free values are bolded). Small world values > 1 indicate non-randomness, all networks were small world. “Best fit” was indicated with the lowest AIC from the gofstat function in the *fitdistrplus* package[2]. The distributions tested with gofstat include: lognormal, poisson, pareto, weibull, and gamma. None of the tested distributions fit the data particularly well (all KS tests rejected, all chi-squared p-values >> 0.001), but the poisson distribution did not perform the best for any network, indicating non-randomness.

| Type | Kingdom | Habitat | Site | Small world value | Power law fit | alpha | "Best fit" distribution from fitdistrplus |
|---|---|---|---|---|---|---|---|
| Watershed | Bacteria | All | All | 10.709 | 0.017 | 10.986 | gamma |
| Watershed | Interkingdom | All | All | 8.841 | 0.000 | 10.693 | gamma |
| Habitat | Interkingdom | Stream | All | 28.283 | 0.002 | 4.710 | weibull |
| Habitat | Bacteria | Terrestrial | All | 21.016 | 0.018 | 7.157 | weibull |
| Habitat | Interkingdom | Terrestrial | All | 23.397 | 0.047 | 7.191 | weibull |

Table S3. Clustering coefficient versus connectance for all non power law networks.

| Type | Kingdom | Habitat | Clustering Coefficient | Connectance |
|---|---|---|---|---|
| Watershed | Bacteria | All | 0.06240 | 0.00240 |
| Watershed | Interkingdom | All | 0.06101 | 0.00294 |
| Habitat | Interkingdom | Stream | 0.17666 | 0.00257 |
| Habitat | Bacteria | Terrestrial | 0.12344 | 0.00244 |
| Habitat | Interkingdom | Terrestrial | 0.13603 | 0.00244 |

Table S4. Pairwise node degree comparisons among bacteria, fungi and interkingdom networks with Tukey HSD test and adjusted P-values for multiple comparisons

| Network Type | Habitat or Site | Comparison | Mean Difference | Adj. P |
|---|---|---|---|---|
| Watershed | All | Fungi-Bacteria | -12.958 | <0.001 |
| Watershed | All | Interkingdom-Bacteria | 7.857 | <0.001 |
| Watershed | All | Interkingdom-Fungi | 20.814 | <0.001 |
| Habitat | Stream | Fungi-Bacteria | -30.395 | <0.001 |
| Habitat | Stream | Interkingdom-Bacteria | 7.572 | <0.001 |
| Habitat | Stream | Interkingdom-Fungi | 37.967 | <0.001 |
| Habitat | Terrestrial | Fungi-Bacteria | -18.178 | <0.001 |
| Habitat | Terrestrial | Interkingdom-Bacteria | 8.181 | <0.001 |
| Habitat | Terrestrial | Interkingdom-Fungi | 26.358 | <0.001 |
| Habitat | Marine | Fungi-Bacteria | -22.170 | <0.001 |
| Habitat | Marine | Interkingdom-Bacteria | 7.265 | <0.001 |
| Habitat | Marine | Interkingdom-Fungi | 29.435 | <0.001 |
| Gradient | Beach | Fungi-Bacteria | -13.618 | <0.001 |
| Gradient | Beach | Interkingdom-Bacteria | 4.531 | <0.001 |
| Gradient | Beach | Interkingdom-Fungi | 18.149 | <0.001 |
| Gradient | Estuary | Fungi-Bacteria | -16.605 | <0.001 |
| Gradient | Estuary | Interkingdom-Bacteria | 10.080 | <0.001 |
| Gradient | Estuary | Interkingdom-Fungi | 26.684 | <0.001 |

| | | | | |
|---|---|---|---|---|
| Gradient | Entrance | Fungi-Bacteria | -22.351 | <0.001 |
| Gradient | Entrance | Interkingdom-Bacteria | 9.597 | <0.001 |
| Gradient | Entrance | Interkingdom-Fungi | 31.948 | <0.001 |
| Gradient | Confluence | Fungi-Bacteria | -20.008 | <0.001 |
| Gradient | Confluence | Interkingdom-Bacteria | 8.831 | <0.001 |
| Gradient | Confluence | Interkingdom-Fungi | 28.839 | <0.001 |
| Gradient | Waterfall | Fungi-Bacteria | -21.522 | <0.001 |
| Gradient | Waterfall | Interkingdom-Bacteria | 6.212 | <0.001 |
| Gradient | Waterfall | Interkingdom-Fungi | 27.734 | <0.001 |
| Gradient | Drum Road | Fungi-Bacteria | -21.510 | <0.001 |
| Gradient | Drum Road | Interkingdom-Bacteria | 8.758 | <0.001 |
| Gradient | Drum Road | Interkingdom-Fungi | 30.268 | <0.001 |
| Gradient | Ridge | Fungi-Bacteria | -14.443 | <0.001 |
| Gradient | Ridge | Interkingdom-Bacteria | 4.052 | <0.001 |
| Gradient | Ridge | Interkingdom-Fungi | 18.495 | <0.001 |

Table S5. Pairwise node betweenness centrality comparisons among bacteria, fungi and interkingdom networks with Tukey HSD test and adjusted P-values for multiple comparisons

| Network Type | Habitat or Site | Comparison | Mean Difference | Adj. P |
|---|---|---|---|---|
| Watershed | All | Fungi-Bacteria | -39.656 | 0.940 |
| Watershed | All | Interkingdom-Bacteria | 642.802 | <0.001 |
| Watershed | All | Interkingdom-Fungi | 682.458 | <0.001 |
| Habitat | Stream | Fungi-Bacteria | -6147.504 | <0.001 |
| Habitat | Stream | Interkingdom-Bacteria | 1523.513 | <0.001 |
| Habitat | Stream | Interkingdom-Fungi | 7671.017 | <0.001 |
| Habitat | Terrestrial | Fungi-Bacteria | -4205.034 | <0.001 |
| Habitat | Terrestrial | Interkingdom-Bacteria | 1535.551 | <0.001 |
| Habitat | Terrestrial | Interkingdom-Fungi | 5740.585 | <0.001 |
| Habitat | Marine | Fungi-Bacteria | -6237.274 | <0.001 |
| Habitat | Marine | Interkingdom-Bacteria | 888.391 | <0.001 |

| | | | | |
|---|---|---|---|---|
| Habitat | Marine | Interkingdom-Fungi | 7125.665 | <0.001 |
| Gradient | Beach | Fungi-Bacteria | -1403.206 | <0.001 |
| Gradient | Beach | Interkingdom-Bacteria | 1122.157 | <0.001 |
| Gradient | Beach | Interkingdom-Fungi | 2525.363 | <0.001 |
| Gradient | Estuary | Fungi-Bacteria | -4158.123 | <0.001 |
| Gradient | Estuary | Interkingdom-Bacteria | 1132.662 | <0.001 |
| Gradient | Estuary | Interkingdom-Fungi | 5290.784 | <0.001 |
| Gradient | Entrance | Fungi-Bacteria | -1827.707 | <0.001 |
| Gradient | Entrance | Interkingdom-Bacteria | 1278.025 | <0.001 |
| Gradient | Entrance | Interkingdom-Fungi | 3105.731 | <0.001 |
| Gradient | Confluence | Fungi-Bacteria | -2171.764 | <0.001 |
| Gradient | Confluence | Interkingdom-Bacteria | 1210.819 | <0.001 |
| Gradient | Confluence | Interkingdom-Fungi | 3382.583 | <0.001 |
| Gradient | Waterfall | Fungi-Bacteria | -2452.087 | <0.001 |
| Gradient | Waterfall | Interkingdom-Bacteria | 1170.632 | <0.001 |
| Gradient | Waterfall | Interkingdom-Fungi | 3622.719 | <0.001 |
| Gradient | Drum Road | Fungi-Bacteria | -2360.065 | <0.001 |
| Gradient | Drum Road | Interkingdom-Bacteria | 1070.018 | <0.001 |
| Gradient | Drum Road | Interkingdom-Fungi | 3430.083 | <0.001 |
| Gradient | Ridge | Fungi-Bacteria | -2207.699 | <0.001 |
| Gradient | Ridge | Interkingdom-Bacteria | 698.594 | 0.033 |
| Gradient | Ridge | Interkingdom-Fungi | 2906.293 | <0.001 |

Table S6. Pairwise node degree comparisons among habitats and site networks with Tukey HSD test and adjusted P-values for multiple comparisons.

| Network Type | Kingdom | Comparison | Mean Difference | Adj. P |
|---|---|---|---|---|
| Habitat | Fungi | Stream-Marine | 3.309 | <0.001 |
| Habitat | Fungi | Terrestrial-Marine | 3.072 | <0.001 |
| Habitat | Fungi | Terrestrial-Stream | -0.237 | 0.271 |
| Habitat | Bacteria | Stream-Marine | 11.534 | <0.001 |

| | | | | |
|---|---|---|---|---|
| Habitat | Bacteria | Terrestrial-Marine | -0.920 | 0.013 |
| Habitat | Bacteria | Terrestrial-Stream | -12.454 | <0.001 |
| Habitat | Interkingdom | Stream-Marine | 11.841 | <0.001 |
| Habitat | Interkingdom | Terrestrial-Marine | -0.004 | 1.000 |
| Habitat | Interkingdom | Terrestrial-Stream | -11.845 | <0.001 |
| Gradient | Fungi | Confluence-Beach | -0.179 | 0.349 |
| Gradient | Fungi | DrumRoad-Beach | -0.038 | 0.999 |
| Gradient | Fungi | Entrance-Beach | -0.159 | 0.485 |
| Gradient | Fungi | Estuary-Beach | 3.544 | <0.001 |
| Gradient | Fungi | Ridge-Beach | -0.807 | <0.001 |
| Gradient | Fungi | Waterfall-Beach | -0.110 | 0.864 |
| Gradient | Fungi | DrumRoad-Confluence | 0.141 | 0.576 |
| Gradient | Fungi | Entrance-Confluence | 0.020 | 1.000 |
| Gradient | Fungi | Estuary-Confluence | 3.723 | <0.001 |
| Gradient | Fungi | Ridge-Confluence | -0.629 | <0.001 |
| Gradient | Fungi | Waterfall-Confluence | 0.068 | 0.980 |
| Gradient | Fungi | Entrance-DrumRoad | -0.121 | 0.726 |
| Gradient | Fungi | Estuary-DrumRoad | 3.582 | <0.001 |
| Gradient | Fungi | Ridge-DrumRoad | -0.770 | <0.001 |
| Gradient | Fungi | Waterfall-DrumRoad | -0.073 | 0.975 |
| Gradient | Fungi | Estuary-Entrance | 3.703 | <0.001 |
| Gradient | Fungi | Ridge-Entrance | -0.648 | <0.001 |
| Gradient | Fungi | Waterfall-Entrance | 0.049 | 0.997 |
| Gradient | Fungi | Ridge-Estuary | -4.351 | <0.001 |
| Gradient | Fungi | Waterfall-Estuary | -3.654 | <0.001 |
| Gradient | Fungi | Waterfall-Ridge | 0.697 | <0.001 |
| Gradient | Bacteria | Confluence-Beach | 6.211 | <0.001 |
| Gradient | Bacteria | DrumRoad-Beach | 7.854 | <0.001 |
| Gradient | Bacteria | Entrance-Beach | 8.574 | <0.001 |
| Gradient | Bacteria | Estuary-Beach | 6.530 | <0.001 |
| Gradient | Bacteria | Ridge-Beach | 0.018 | 1.000 |
| Gradient | Bacteria | Waterfall-Beach | 7.793 | <0.001 |

| | | | | |
|---|---|---|---|---|
| Gradient | Bacteria | DrumRoad-Confluence | 1.643 | <0.001 |
| Gradient | Bacteria | Entrance-Confluence | 2.363 | <0.001 |
| Gradient | Bacteria | Estuary-Confluence | 0.320 | 0.879 |
| Gradient | Bacteria | Ridge-Confluence | -6.193 | <0.001 |
| Gradient | Bacteria | Waterfall-Confluence | 1.583 | <0.001 |
| Gradient | Bacteria | Entrance-DrumRoad | 0.720 | 0.062 |
| Gradient | Bacteria | Estuary-DrumRoad | -1.324 | <0.001 |
| Gradient | Bacteria | Ridge-DrumRoad | -7.836 | <0.001 |
| Gradient | Bacteria | Waterfall-DrumRoad | -0.061 | 1.000 |
| Gradient | Bacteria | Estuary-Entrance | -2.044 | <0.001 |
| Gradient | Bacteria | Ridge-Entrance | -8.556 | <0.001 |
| Gradient | Bacteria | Waterfall-Entrance | -0.781 | 0.033 |
| Gradient | Bacteria | Ridge-Estuary | -6.513 | <0.001 |
| Gradient | Bacteria | Waterfall-Estuary | 1.263 | <0.001 |
| Gradient | Bacteria | Waterfall-Ridge | 7.776 | <0.001 |
| Gradient | Interkingdom | Confluence-Beach | 10.511 | <0.001 |
| Gradient | Interkingdom | DrumRoad-Beach | 12.081 | <0.001 |
| Gradient | Interkingdom | Entrance-Beach | 13.640 | <0.001 |
| Gradient | Interkingdom | Estuary-Beach | 12.079 | <0.001 |
| Gradient | Interkingdom | Ridge-Beach | -0.461 | 0.792 |
| Gradient | Interkingdom | Waterfall-Beach | 9.475 | <0.001 |
| Gradient | Interkingdom | DrumRoad-Confluence | 1.570 | <0.001 |
| Gradient | Interkingdom | Entrance-Confluence | 3.129 | <0.001 |
| Gradient | Interkingdom | Estuary-Confluence | 1.568 | <0.001 |
| Gradient | Interkingdom | Ridge-Confluence | -10.973 | <0.001 |
| Gradient | Interkingdom | Waterfall-Confluence | -1.037 | 0.004 |
| Gradient | Interkingdom | Entrance-DrumRoad | 1.559 | <0.001 |
| Gradient | Interkingdom | Estuary-DrumRoad | -0.002 | 1.000 |
| Gradient | Interkingdom | Ridge-DrumRoad | -12.542 | <0.001 |
| Gradient | Interkingdom | Waterfall-DrumRoad | -2.606 | <0.001 |
| Gradient | Interkingdom | Estuary-Entrance | -1.561 | <0.001 |
| Gradient | Interkingdom | Ridge-Entrance | -14.101 | <0.001 |

| | | | | |
|---|---|---|---|---|
| Gradient | Interkingdom | Waterfall-Entrance | -4.165 | <0.001 |
| Gradient | Interkingdom | Ridge-Estuary | -12.541 | <0.001 |
| Gradient | Interkingdom | Waterfall-Estuary | -2.605 | <0.001 |
| Gradient | Interkingdom | Waterfall-Ridge | 9.936 | <0.001 |

Table S7. Pairwise node betweenness centrality comparisons among habitats and site networks with Tukey HSD test and adjusted P-values for multiple comparisons.

| Network Type | Kingdom | Comparison | Mean Difference | Adj. P |
|---|---|---|---|---|
| Habitat | Fungi | Stream-Marine | 199.155 | 0.101 |
| Habitat | Fungi | Terrestrial-Marine | 167.443 | 0.204 |
| Habitat | Fungi | Terrestrial-Stream | -31.712 | 0.920 |
| Habitat | Bacteria | Stream-Marine | 109.385 | 0.823 |
| Habitat | Bacteria | Terrestrial-Marine | -1864.797 | <0.001 |
| Habitat | Bacteria | Terrestrial-Stream | -1974.182 | <0.001 |
| Habitat | Interkingdom | Stream-Marine | 744.507 | <0.001 |
| Habitat | Interkingdom | Terrestrial-Marine | -1217.636 | <0.001 |
| Habitat | Interkingdom | Terrestrial-Stream | -1962.143 | <0.001 |
| Gradient | Fungi | Confluence-Beach | 1040.898 | <0.001 |
| Gradient | Fungi | DrumRoad-Beach | 732.050 | 0.011 |
| Gradient | Fungi | Entrance-Beach | 1462.376 | <0.001 |
| Gradient | Fungi | Estuary-Beach | -755.988 | 0.005 |
| Gradient | Fungi | Ridge-Beach | -185.904 | 0.990 |
| Gradient | Fungi | Waterfall-Beach | 701.863 | 0.018 |
| Gradient | Fungi | DrumRoad-Confluence | -308.848 | 0.707 |
| Gradient | Fungi | Entrance-Confluence | 421.479 | 0.300 |
| Gradient | Fungi | Estuary-Confluence | -1796.886 | <0.001 |
| Gradient | Fungi | Ridge-Confluence | -1226.801 | <0.001 |
| Gradient | Fungi | Waterfall-Confluence | -339.034 | 0.614 |
| Gradient | Fungi | Entrance-DrumRoad | 730.326 | 0.004 |
| Gradient | Fungi | Estuary-DrumRoad | -1488.038 | <0.001 |

| | | | | |
|---|---|---|---|---|
| Gradient | Fungi | Ridge-DrumRoad | -917.953 | 0.002 |
| Gradient | Fungi | Waterfall-DrumRoad | -30.187 | 1.000 |
| Gradient | Fungi | Estuary-Entrance | -2218.364 | <0.001 |
| Gradient | Fungi | Ridge-Entrance | -1648.280 | <0.001 |
| Gradient | Fungi | Waterfall-Entrance | -760.513 | 0.002 |
| Gradient | Fungi | Ridge-Estuary | 570.084 | 0.188 |
| Gradient | Fungi | Waterfall-Estuary | 1457.851 | <0.001 |
| Gradient | Fungi | Waterfall-Ridge | 887.767 | 0.004 |
| Gradient | Bacteria | Confluence-Beach | 1809.456 | <0.001 |
| Gradient | Bacteria | DrumRoad-Beach | 1688.909 | <0.001 |
| Gradient | Bacteria | Entrance-Beach | 1886.878 | <0.001 |
| Gradient | Bacteria | Estuary-Beach | 1998.929 | <0.001 |
| Gradient | Bacteria | Ridge-Beach | 618.590 | 0.048 |
| Gradient | Bacteria | Waterfall-Beach | 1750.745 | <0.001 |
| Gradient | Bacteria | DrumRoad-Confluence | -120.547 | 0.994 |
| Gradient | Bacteria | Entrance-Confluence | 77.421 | 0.999 |
| Gradient | Bacteria | Estuary-Confluence | 189.473 | 0.941 |
| Gradient | Bacteria | Ridge-Confluence | -1190.867 | <0.001 |
| Gradient | Bacteria | Waterfall-Confluence | -58.711 | 1.000 |
| Gradient | Bacteria | Entrance-DrumRoad | 197.968 | 0.917 |
| Gradient | Bacteria | Estuary-DrumRoad | 310.020 | 0.591 |
| Gradient | Bacteria | Ridge-DrumRoad | -1070.320 | <0.001 |
| Gradient | Bacteria | Waterfall-DrumRoad | 61.836 | 1.000 |
| Gradient | Bacteria | Estuary-Entrance | 112.051 | 0.995 |
| Gradient | Bacteria | Ridge-Entrance | -1268.288 | <0.001 |
| Gradient | Bacteria | Waterfall-Entrance | -136.132 | 0.987 |
| Gradient | Bacteria | Ridge-Estuary | -1380.339 | <0.001 |
| Gradient | Bacteria | Waterfall-Estuary | -248.184 | 0.813 |
| Gradient | Bacteria | Waterfall-Ridge | 1132.156 | <0.001 |
| Gradient | Interkingdom | Confluence-Beach | 1898.118 | <0.001 |
| Gradient | Interkingdom | DrumRoad-Beach | 1636.770 | <0.001 |
| Gradient | Interkingdom | Entrance-Beach | 2042.745 | <0.001 |

| | | | | |
|---|---|---|---|---|
| Gradient | Interkingdom | Estuary-Beach | 2009.433 | <0.001 |
| Gradient | Interkingdom | Ridge-Beach | 195.027 | 0.962 |
| Gradient | Interkingdom | Waterfall-Beach | 1799.220 | <0.001 |
| Gradient | Interkingdom | DrumRoad-Confluence | -261.348 | 0.732 |
| Gradient | Interkingdom | Entrance-Confluence | 144.627 | 0.978 |
| Gradient | Interkingdom | Estuary-Confluence | 111.316 | 0.995 |
| Gradient | Interkingdom | Ridge-Confluence | -1703.091 | <0.001 |
| Gradient | Interkingdom | Waterfall-Confluence | -98.898 | 0.998 |
| Gradient | Interkingdom | Entrance-DrumRoad | 405.975 | 0.191 |
| Gradient | Interkingdom | Estuary-DrumRoad | 372.664 | 0.308 |
| Gradient | Interkingdom | Ridge-DrumRoad | -1441.743 | <0.001 |
| Gradient | Interkingdom | Waterfall-DrumRoad | 162.450 | 0.966 |
| Gradient | Interkingdom | Estuary-Entrance | -33.311 | 1.000 |
| Gradient | Interkingdom | Ridge-Entrance | -1847.718 | <0.001 |
| Gradient | Interkingdom | Waterfall-Entrance | -243.525 | 0.780 |
| Gradient | Interkingdom | Ridge-Estuary | -1814.407 | <0.001 |
| Gradient | Interkingdom | Waterfall-Estuary | -210.214 | 0.887 |
| Gradient | Interkingdom | Waterfall-Ridge | 1604.193 | <0.001 |

Table S8. Area under the curve (AUC) values for all networks, full and downsampled to the smallest full network (721 nodes).

| | | Robustness | |
|---|---|---|---|
| Type | Kingdom | Full | Downsampled |
| Watershed | Fungi | 0.050 | 0.037 |
| Watershed | Bacteria | 0.201 | 0.195 |
| Watershed | Interkingdom | 0.229 | 0.263 |
| Marine | Fungi | 0.087 | 0.088 |
| Marine | Bacteria | 0.236 | 0.113 |
| Marine | Interkingdom | 0.225 | 0.082 |
| Stream | Fungi | 0.287 | 0.173 |

| | | | |
|---|---|---|---|
| Stream | Bacteria | 0.272 | 0.171 |
| Stream | Interkingdom | 0.298 | 0.187 |
| Terrestrial | Fungi | 0.241 | 0.164 |
| Terrestrial | Bacteria | 0.276 | 0.199 |
| Terrestrial | Interkingdom | 0.292 | 0.181 |
| Beach | Fungi | 0.073 | 0.038 |
| Beach | Bacteria | 0.206 | 0.163 |
| Beach | Interkingdom | 0.222 | 0.137 |
| Estuary | Fungi | 0.194 | 0.153 |
| Estuary | Bacteria | 0.202 | 0.164 |
| Estuary | Interkingdom | 0.217 | 0.155 |
| Entrance | Fungi | 0.066 | 0.021 |
| Entrance | Bacteria | 0.274 | 0.203 |
| Entrance | Interkingdom | 0.276 | 0.174 |
| Confluence | Fungi | 0.071 | 0.020 |
| Confluence | Bacteria | 0.205 | 0.185 |
| Confluence | Interkingdom | 0.232 | 0.174 |
| Waterfall | Fungi | 0.068 | 0.019 |
| Waterfall | Bacteria | 0.204 | 0.157 |
| Waterfall | Interkingdom | 0.222 | 0.147 |
| DrumRoad | Fungi | 0.078 | 0.022 |
| DrumRoad | Bacteria | 0.211 | 0.172 |
| DrumRoad | Interkingdom | 0.230 | 0.172 |
| Ridge | Fungi | 0.038 | 0.038 |
| Ridge | Bacteria | 0.094 | 0.109 |
| Ridge | Interkingdom | 0.119 | 0.141 |

Table S9. Pairwise comparisons of the standardized effect scores of mean phylogenetic distances between bacteria of different habitats. Values in bold indicate significant differences between habitats, as determined by ANOVA and Tukey’s post hoc tests.

| Habitat | Mean Difference | P |
|---|---|---|
| Riverine-Marine | 1.230 | <**0.001** |

| | | |
|---|---|---|
| Terrestrial-Marine | 0.884 | **<0.001** |
| Terrestrial-Riverine | -0.346 | **0.042** |

Table 10. Pairwise comparisons of the standardized effect scores of mean phylogenetic distances between bacteria of different sites along the terrestrial gradient. Values in bold indicate significant differences between sites, as determined by ANOVA and Tukey’s post hoc tests.

| Site | Mean difference | P |
|---|---|---|
| Estuary-Beach | 0.360 | 0.620 |
| Entrance-Beach | -0.313 | 0.761 |
| Confluence-Beach | -0.001 | 1.000 |
| Waterfall-Beach | -0.061 | 1.000 |
| DrumRoad-Beach | -0.277 | 0.851 |
| Ridge-Beach | -1.081 | **<0.001** |
| Entrance-Estuary | -0.673 | **0.028** |
| Confluence-Estuary | -0.362 | 0.616 |
| Waterfall-Estuary | -0.421 | 0.428 |
| Drum Road-Estuary | -0.637 | **0.046** |
| Ridge-Estuary | -1.441 | **<0.001** |
| Confluence-Entrance | 0.312 | 0.765 |
| Waterfall-Entrance | 0.252 | 0.900 |
| Drum Road-Entrance | 0.036 | 1.000 |
| Ridge-Entrance | -0.768 | **0.007** |
| Waterfall-Confluence | -0.060 | 1.000 |
| Drum Road-Confluence | -0.275 | 0.854 |
| Ridge-Confluence | -1.079 | **<0.001** |
| Drum Road-Waterfall | -0.216 | 0.951 |
| Ridge-Waterfall | -1.019 | **<0.001** |
| Ridge-Drum Road | -0.804 | **0.004** |
